# The effect of habitat choice on evolutionary rescue in subdivided populations

**DOI:** 10.1101/738898

**Authors:** Peter Czuppon, François Blanquart, Hildegard Uecker, Florence Débarre

**Affiliations:** Center for Interdisciplinary Research in Biology, CNRS, Collège de France, PSL Research University, Paris, France; Institute of Ecology and Environmental Sciences of Paris, Sorbonne Université, UPEC, CNRS, IRD, INRA, Paris, France; IAME, UMR 1137, INSERM, Université Paris Diderot, Paris, France; Research Group “Stochastic evolutionary dynamics”, Department of Evolutionary Theory, Max-Planck Institute for Evolutionary Biology, Plön, Germany

**Keywords:** evolutionary rescue, local adaptation, source-sink dynamics, dispersal, gene flow, habitat choice, density-dependent dispersal

## Abstract

Evolutionary rescue is the process by which a population, in response to an environmental change, successfully avoids extinction through adaptation. In spatially structured environments, dispersal can affect the probability of rescue. Here, we model an environment consisting of patches that degrade one after another, and we investigate the probability of rescue by a mutant adapted to the degraded habitat. We focus on the effects of dispersal and of immigration biases. We find that the probability of evolutionary rescue can undergo up to three phases: (i) starting from low dispersal rates, it increases with dispersal; (ii) at intermediate dispersal rates, it decreases; (iii) finally, at large dispersal rates, the probability of rescue increases again with dispersal, except if mutants are too counter-selected in not-yet-degraded patches. The probability of rescue is generally highest when mutant and wild-type individuals preferentially immigrate into patches that have already undergone environmental change. Additionally, we find that mutants that will eventually rescue the population most likely first appear in non-degraded patches, and that the relative contribution of standing genetic variation vs. de-novo mutations declines with increasing emigration rates. Overall, our results show that habitat choice, when compared to the often studied unbiased immigration scheme, can substantially alter the dynamics of population survival and adaptation to new environments.

## 1. Introduction

Current anthropogenic environmental changes such as deforestation, soil and water contamination or rising temperatures, contribute to the decline of the populations of many species, that might eventually go extinct (Diniz-Filho et al., 2019). Pests and pathogens experience similarly strong selective pressures as a result of consumption of antibiotics and use of pesticides (Ramsayer et al., 2013; Kreiner et al., 2018). The process of genetic adaptation that saves populations from extinction is termed evolutionary rescue. This process is characterized by an initial population decline (that, without adaptation, would result in population extinction) followed by recovery due to the establishment of adapted genotypes, classically resulting in a U-shaped demographic trajectory over time (Gomulkiewicz and Holt, 1995). In recent years, empirical examples of evolutionary rescue have accumulated (as reviewed by Alexander et al., 2014; Carlson et al., 2014; Bell, 2017). Laboratory experiments have provided direct evidence of evolutionary rescue (e.g. Bell and Gonzalez, 2009; Agashe et al., 2011; Lachapelle and Bell, 2012; Lindsey et al., 2013; Stelkens et al., 2014). In the wild, however, demographic and genotypic data are rarely monitored together at the same time, which impedes direct observation of evolutionary rescue. Still, evolutionary rescue has been suggested as a mechanism that has saved a few wild populations from extinction (e.g. Vander Wal et al., 2012; Di Giallonardo and Holmes, 2015; Gignoux-Wolfsohn et al., 2018).

Here, we study the effect of dispersal and habitat choice on evolutionary rescue in a subdivided population. We assume that dispersal intensity and habitat choice are fixed and do not evolve. In general, the traits involved in adaptation can be continuous (e.g. Bürger and Lynch, 1995; Gomulkiewicz and Holt, 1995; Boulding and Hay, 2001; Osmond et al., 2017) or discrete (e.g. Orr and Unckless, 2008; Martin et al., 2013; Uecker et al., 2014). We consider genetic adaptation mediated by a discrete trait and we assume that fitness of individuals is determined by a single haploid locus.

Evolutionary rescue is often studied in a spatially homogeneous situation where the whole population experiences a sudden decrease in habitat quality. In this setting, a large number of theoretical results have been established, for example on the effects of recombination (Uecker and Hermisson, 2016) and horizontal gene transfer (Tazzyman and Bonhoeffer, 2014), mode of reproduction (Glémin and Ronfort, 2013; Uecker, 2017), intra- and interspecific competition (Osmond and de Mazancourt, 2013), predation pressure (Yamamichi and Miner, 2015), bottlenecks (Martin et al., 2013), different genetic pathways (Osmond et al., 2019), and the context-dependent fitness effects of mutations (Anciaux et al., 2018). In contrast to these abrupt change scenarios, evolutionary rescue can also be studied in a gradually changing environment (e.g. Bürger and Lynch, 1995; Osmond et al., 2017).

In fragmented environments, habitat deterioration is not necessarily synchronized across patches: there can be a transient spatially heterogeneous environment consisting of a mosaic of old- and of degraded-habitat patches, until eventually the whole environment has deteriorated. If individuals that populate different patches are able to move between those, the effect of dispersal on evolutionary rescue needs to be taken into account (Uecker et al., 2014; Tomasini and Peischl, 2020). The intensity of dispersal among patches tunes how abruptly environmental change is experienced. With very low dispersal, patches are essentially isolated from each other, and each patch undergoes an abrupt change independently of the other patches. With higher dispersal, asynchronous deterioration among patches is experienced as a more gradual change overall. Experiments that study the effect of dispersal on evolutionary rescue are rare, but, for instance, Bell and Gonzalez (2011) found that dispersal can increase the probability of successful genetic adaptation.

The transient mosaic of degraded and non-degraded patches that results from asynchronous degradation in a fragmented habitat is similar to the setting of models of source-sink dynamics. These models represent a spatially heterogeneous environment, constant in time, in which wild-type populations in unfavorable habitats can only be maintained thanks to recurrent immigration from favorable habitats. Experimental and theoretical studies have found that an increase in dispersal can have a positive or a negative effect on genetic adaptation in a heterogeneous environment (see e.g., for studies on discrete traits, Holt and Gomulkiewicz (1997); Gomulkiewicz et al. (1999) for positive effects; Nagylaki (1978); Karlin and Campbell (1981); Storfer and Sih (1998) for negative effects; and Kawecki (2000); Gallet et al. (2018) for both effects).

In theoretical studies of local adaptation and evolutionary rescue, dispersal is typically assumed to be unbiased, i.e. dispersing individuals are distributed uniformly among patches. Only few investigations in the context of local adaptation in source-sink systems have taken into account non-uniform dispersal patterns (e.g. Kawecki, 1995; Holt, 1996; Kawecki and Holt, 2002; Amarasekare, 2004). This analytical focus on unbiased dispersal is in stark contrast to dispersal schemes observed in nature (Edelaar et al., 2008; Clobert et al., 2009; Edelaar and Bolnick, 2012).

One of the best documented modes of non-uniform dispersal is density-dependent dispersal. Density dependence can be positive or negative: either individuals prefer to settle or stay in large groups (positive density-dependence), or they choose to remain in or move to less populated regions (negative density-dependence). Density-dependent dispersal, of either form, is ubiquitously found in nature and has been reported in many species across the tree of life, including insects (Endriss et al., 2019), spiders (De Meester and Bonte, 2010), amphibians (Gautier et al., 2006), birds (Wilson et al., 2017b), fishes (Turgeon and Kramer, 2012), and mammals (Støen et al., 2006).

Another well-established dispersal scheme is a type of habitat choice whereby individuals tend to immigrate into habitats to which they are best adapted. This mechanism has for example been reported in lizards (Bestion et al., 2015), birds (Dreiss et al., 2011; Benkman, 2017), fishes (Bolnick et al., 2009), worms (Mathieu et al., 2010), and ciliates (Jacob et al., 2017, 2018).

Dispersal biases can affect the different steps of dispersal (the probability to emigrate, the vagrant stage, and immigration (Bowler and Benton, 2005; Ronce, 2007)). In this work, we focus on effects on the immigration step.

We model an environment that consists of various patches with one of two possible habitats: the ‘old’ habitat, in which both types, wild type and mutant, have sufficient offspring (on average) to ensure survival of the population, and the degraded ‘new’ habitat, where in the absence of immigration the wild-type population will eventually go extinct. We study four biologically motivated dispersal schemes, which correspond to the four combinations of biases towards old vs. new patches for wild type and mutants, and we compare these dispersal schemes to unbiased dispersal. Our analysis is carried out step-wise. We first consider a temporally constant but spatially heterogeneous environment with two (‘old’ and ‘new’) patch types. In this setting, we first study the probability of establishment of a single mutant, assuming there are no further mutations between types. We then relax the assumption of no further mutations, and compute a probability of adaptation, i.e. of emergence and successful establishment of the mutant lineage. Finally, we let habitat degradation proceed, assuming that patches, one after another, deteriorate over time until all locations contain the new habitat. Using the previous results, we approximate the probability of evolutionary rescue, i.e. that a mutation appears and establishes, thereby allowing the population to persist in spite of environmental degradation. We find that dispersal biases affect the probabilities of establishment and of evolutionary rescue.

## 2. Model and methods

### 2.1. Main assumptions and life-cycle

We consider a spatially structured environment consisting of *M* patches all connected to each other. The habitat of a patch is either in the *old* or in the *new* state, corresponding to habitat quality before and after environmental deterioration, respectively. One after another every *τ* generations, the habitat of a patch deteriorates, from old to new state, the transition being irreversible. Initially (time *t* < 0), all patches are of the old-habitat type. At time *t* = 0, the first patch deteriorates. After (*M* − 1)*τ* generations, all patches are of the new-habitat type. We denote the time-dependent frequency of old-habitat patches by *f*_old_. It equals 1 before the first environmental change takes place (*t* < 0), and decreases by 1/*M* after each environmental deterioration event, until it eventually hits 0, when all patches have undergone the environmental change. This setting corresponds to the one analyzed by Uecker et al. (2014), and more recently by Tomasini and Peischl (2020) in the special case of just two patches. The maximum numbers of individuals that can live in a patch of a given habitat type, i.e. the carrying capacities, are denoted *K*_old_ and *K*_new_ for old- and new-habitat patches respectively; *K*_old_ and *K*_new_ may differ. We view these carrying capacities as a number of territories or nesting sites; all of these sites are assumed to be accessible to individuals of both types, so that *K*_old_ and *K*_new_ are the same for both types of individuals.

The population living in this environment consists of asexually reproducing, haploid individuals; generations are discrete and non-overlapping. There are two possible types of individuals, wild types and mutants. The individuals go through the following life-cycle:

i. Dispersal: individuals may move between patches. Further details about this step are given below.
ii. Reproduction: individuals reproduce within patches. The number of offspring produced by an individual of type *i* (before density regulation, if any), i.e. its fecundity, is drawn from a Poisson distribution with expectation 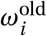 and 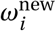 in old- and new-habitat patches, respectively. Having fewer than 1 offspring in expectation means that the local subpopulation will get extinct in the absence of immigration, because the deaths of the parents at each generation are not compensated by enough births on average. On the contrary, the local population is viable, i.e. has a chance not to go extinct, if the expected fecundity is greater than 1. In most figures, we assume that both wild-type and mutant individuals have an expected fecundity greater than 1 in old-habitat patches, and that the mutant’s expected fecundity there is lower than the wild type’s, 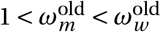—but we also consider the extreme scenario 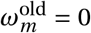. In new-habitat patches, a wild-type population will eventually go extinct, while a mutant one would persist, hence the term “rescue mutant”: 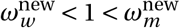. All parents die at the end of this step.
iii. Mutation: wild-type offspring mutate to the rescue mutant type with probability *θ*. Back mutations from the mutant to the wild type are neglected.
iv. Density regulation: if the number of offspring produced locally exceeds the local carrying capacity *K_k_* (where *k* refers to the habitat type, old or new), the population size is down-regulated to *K_k_* by randomly removing individuals until the local population size is equal to *K_k_* (ceiling density regulation). Mutant offspring have the same chance of being removed as wild-type offspring, i.e. we assume that wild-type and mutant individuals are competitively equivalent. If the number of offspring is below the carrying capacity, the regulation step is unnecessary. We write “successful offspring” for offspring that survive the density regulation step, and become adults at the next generation. At the end of this step, all offspring become adults, and a new cycle begins.

### 2.2. Dispersal mechanisms

We split the dispersal step into emigration and immigration. Emigration is type-independent: all individuals have the same probability *m* of leaving the patch they were born in. We assume that dispersal biases affect the immigration step. We denote by *π_i_* the bias for immigration to an old-habitat patch, where the index *i* refers to the type of the dispersing individual (*w* for wild type, *m* for mutant). If *π_i_* < 0, individuals of type *i* are relatively more likely to settle in new-habitat patches than in old-habitat patches; conversely, their bias is towards old-habitat patches if *π_i_ >* 0. The case *π_i_* = 0 corresponds to unbiased dispersal. For simplicity, we assume that dispersal is cost-free. While local population sizes may be affected by dispersal, the global size of the metapopulation remains the same before and after dispersal. Note that our methods can readily be applied to costly dispersal (including to costs that differ among wild-type and mutant individuals), and also to type- and habitat-dependent emigration probabilities.

The probability that a dispersing individual of type *i* settles in a patch of the new-habitat type is

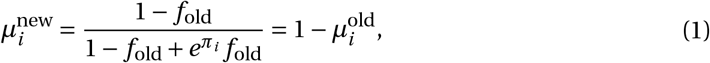

where, as defined above, *f*_old_, is the frequency of old-habitat patches and *π_i_* the dispersal bias into old-habitat patches. The use of an exponential *e^π_i_^* ensures that the fraction in Eq. (1) is positive and between zero and one.

Qualitatively, there are four possible combinations of dispersal biases. We name them according to the preferences of wild type and then of the mutant (e.g., “Old-New”, wild-type individuals have a bias toward old-habitat patches, and mutant individuals toward new-habitat patches). We add to these four dispersal schemes the case of unbiased dispersal. Fig. 1 provides an overview of the different schemes, together with the parameter values used in the numerical simulations.

**Figure 1:**
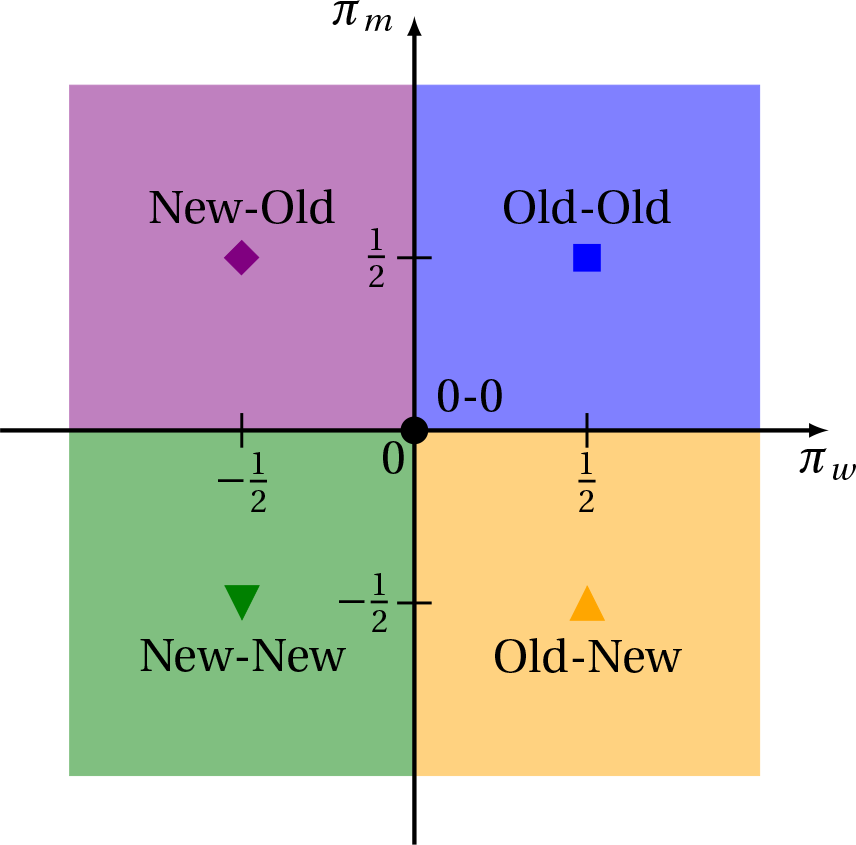
Parameter sets and legends for the different dispersal schemes. The colors and markers are the same across all figures. The horizontal axis is the dispersal bias of the wild type, *π_w_* (positive values corresponding to preferential immigration into old-habitat patches), and the vertical axis that of the mutant, *π_m_*. The markers are located at the parameter values used in the simulations.

Each of these dispersal schemes can be related to a biological illustration:

**Old-Old (*π_w_* > 0, *π_m_* > 0)** Both types of individuals have a bias towards old-habitat patches. If we assume that mutant individuals have a higher fecundity in old-habitat patches than in new-habitat patches (i.e., 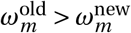, which is the case in our numerical examples), then this dispersal scheme corresponds to biases towards the habitat where individuals have the highest fecundity. This type of dispersal, which can be described as matching habitat choice, has for example been observed with common lizards *Zootoca vivipara* (Bestion et al., 2015), three-spine sticklebacks *Gasterosteus aculeatus* (Bolnick et al., 2009), and barn owls *Tyto alba* (Dreiss et al., 2011). Population densities being high in the old-habitat patches, this dispersal scheme can also be interpreted as positive density-dependent immigration. For prey species, highly populated locations can be an indication for a safe shelter, or of a place with numerous mating opportunities. This type of positive density-dependent immigration (also called conspecific attraction) is for example found in several amphibians, e.g. the salamander species *Mertensiella luschani* (Gautier et al., 2006) and *Ambystoma maculatum* (Greene et al., 2016) or the frogs *Oophaga pumilio* (Folt et al., 2018).
**Old-New (*π_w_* > 0, *π_m_* < 0)** Wild-type individuals preferentially immigrate into old-habitat patches, while mutants prefer new-habitat patches. This corresponds to immigration to patches where the focal type is fitter than the other type (since 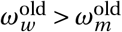 and 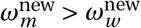). A similar dispersal scheme was recently observed for the ciliates *Tetrahymena thermophila* with a specialist and generalist type (Jacob et al., 2018), and where the specialist disperses to its preferred habitat while the generalist prefers to immigrate to a suboptimal habitat where it outcompetes the specialist.
**New-New (*π_w_* < 0, *π_m_* < 0)** Both types of individuals preferentially immigrate into new-habitat patches. Population densities being on average lower in new-habitat patches, and in particular, because the carrying capacity is not typically reached in new-habitat patches during the initial phase of evolutionary rescue, this dispersal scheme can be interpreted as negative density-dependent immigration, whereby individuals are more likely to move to less populated patches. In nature, such a bias may exist because, in less populated locations, resources might be more abundant, intra-specific competition alleviated and the chance of infection transmission decreased, which may compensate for the potentially reduced habitat quality. Density-dependent immigration effects as described here, are for example found in the damselfish species *Stegastes adustus* (Turgeon and Kramer, 2012) and the migratory birds *Setophaga ruticilla* (Wilson et al., 2017b).
**New-Old (*π_w_* < 0, *π_m_* > 0)** Wild-type individuals preferentially immigrate into new-habitat patches, while mutants prefer old-habitat patches. This dispersal scheme is considered mostly for completeness, because it is biologically quite unlikely. It can be related to the concept of an ‘ecological trap’, wherein individuals tend to immigrate into patches that cannot sustain a population, in its most extreme form resulting in the extinction of the species (Battin, 2004).
**Unbiased dispersal (o-o) (*π_w_* = 0, *π_m_* = 0)** Neither type has a dispersal bias. Most theoretical results examining the interplay of dispersal and establishment have used this dispersal scheme. We therefore use it as a benchmark to which we compare the biased dispersal schemes.

All the model parameters are summarized in Table 1 along with the default parameter values and ranges. If not stated otherwise, the default parameter values are used for the stochastic simulations.

**Table 1:**
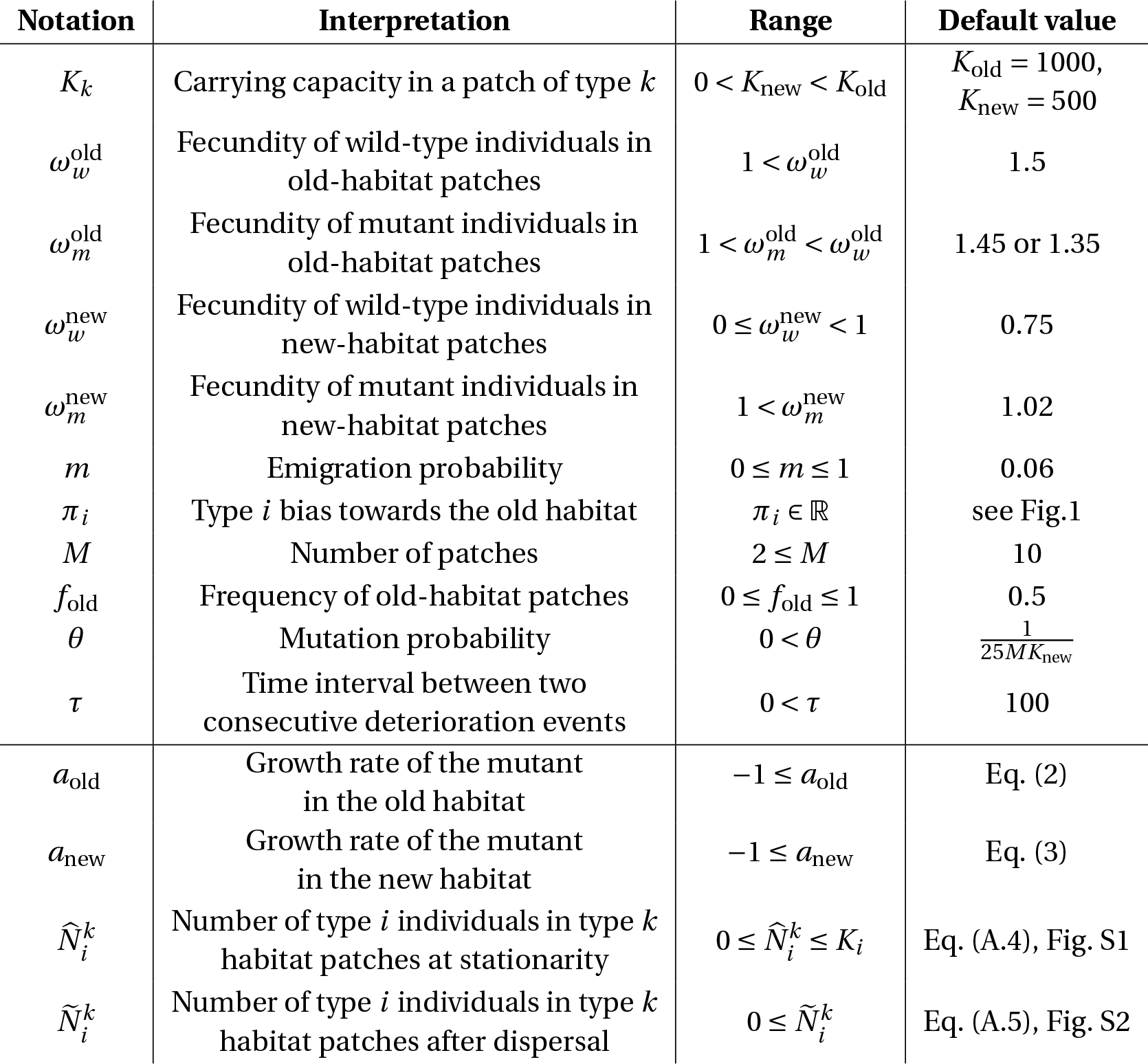
Model parameters and variables.

### 2.3. Analysis steps

We decompose our analysis into several steps of increasing complexity.

1. We first consider an environment that is constant over time, and heterogeneous over space, with a fraction *f*_old_ of old-habitat patches and 1 − *f*_old_ of new-habitat patches. The population is initiated with wild-type individuals at carrying capacity in old-habitat patches, and at the migration-selection equilibrium 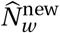 in new-habitat patches (see Section A in the Supplementary Information (SI) for details), and with a single mutant individual, either in an old- or in a new-habitat patch. There are no further mutations (*θ* = 0), and we compute the *probability of establishment* of a mutant lineage.
2. We then consider the same environmental setting, but initialize the population with only wild-type individuals. Mutants can appear by mutation during the simulation (*θ >* 0). We compute the *probability of adaptation*, i.e. that, during a fixed time interval, a mutant appears by mutation and then establishes.
3. Finally, we consider the full scenario where each patch degrades one after another, as described above. The environment is spatially and temporally variable. The population is initialized with only old-habitat patches, all at carrying capacity, with wild-type individuals only. We compute the *probability of evolutionary rescue*, i.e. that a mutant appears by mutation and establishes before the population goes extinct.

### 2.4. Additional assumptions for the analytical part

We make a few additional assumptions in the analytical part of our work; these assumptions are relaxed in the stochastic simulations.

A key assumption to our mathematical analysis is that the mutant individuals are rare enough that their dynamics do not affect the wild-type population during the establishment phase of the mutant lineage. Because of their rarity, we can also consider that all mutants reproduce, disperse and die independently of each other. The wild-type population sets a demographic context that affects mutant dynamics. The mathematical analysis therefore focuses on the population dynamics of the mutant population, considering the wild-type population as constant over time (except in the rescue scenario).

We assume that the subpopulations in old-habitat patches are always at carrying capacity, i.e. that there are always enough offspring that are produced to at least replace all the parents. Denoting by 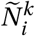 the number of type-*i* individuals in a *k*-habitat patch right after dispersal, then the expected number of successful offspring of mutant individuals in this old-habitat patch (i.e., of offspring that survive density regulation and become adults in the next generation) is

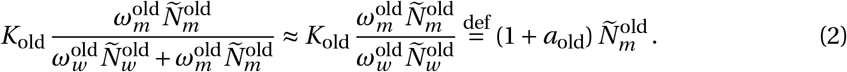

The approximation results from the assumption that mutants are rare compared to wildtype individuals in old-habitat patches. Eq. (2) defines the per-capita expected growth rate of mutants in old-habitat patches, *a*_old_. It depends on 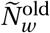, the size of the local wild-type population right after dispersal, which is calculated in Section A of the SI (Eq. (A.5a)).

In new-habitat patches, the situation is a bit more involved. Either the local population size after reproduction exceeds the carrying capacity, in which case density regulation is necessary, or it is below the carrying capacity. In the latter case, the expected number of offspring per mutant is simply given by their fecundity 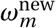. In the former case where the population after reproduction exceeds the carrying capacity *K*_new_, a similar argument as in old-habitat patches allows us to approximate the per capita number of successful offspring as in Eq. (2). These two cases yield the following definition of the per capita growth rate of mutants in new-habitat patches, *a*_new_:

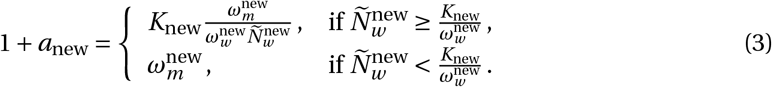

Note, that the first line is obtained using the same rare-mutant approximation as in Eq. (2).

We finally combine the different steps of the life cycle. The expected per capita numbers of successful mutant offspring in habitat *k′* of an individual in a *k*-habitat patch at the beginning of the generation, *λ_k,k′_*, are the following:

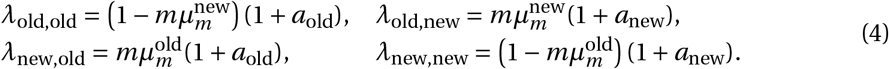

Our final assumption for the mathematical analysis is that the distributions of numbers of successful offspring are Poisson, with means *λ_k,k′_* (counting the successful offspring in habitat *k′* of a parent in a *k*-habitat patch at the beginning of the generation). In reality, only the production of offspring before density regulation is Poisson; here we lump in the effects of dispersal and of density regulation. These mean values are treated as (piecewise-)constant over time. This way, the dynamics of the mutant population can be described by a two-type branching process, for which an established methodology exists (Haccou et al., 2005). (In our context, the two “types” in the name of the method, “two-type branching process”, correspond to the two habitat types.)

All of the assumptions of this section, made for the sake of mathematical analysis, are relaxed in our stochastic simulations.

### 2.5. Simulations

The simulation algorithm implements the life cycle described above. We specify here the sampling distributions that we use.

i. Dispersal: for each patch, a random number of dispersing individuals is drawn from a binomial distribution with success probability *m*. The dispersing individuals from all patches are pooled together and redistributed into patches according to their type and the dispersal pattern. For each type of individuals (wild type and mutant), immigration patches are assigned by first drawing the number of individuals who immigrate into old-habitat patches from a binomial distribution with success probability 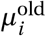 (Eq. (1)), and then distributing these individuals uniformly at random over the old-habitat patches. The remaining dispersing individuals are then distributed uniformly at random into the new-habitat patches.
ii. Reproduction: in each patch, reproduction is simulated by drawing a Poisson distributed number of offspring for each type. The mean of the Poisson distribution is the product of the number of individuals of type *i* in that patch times 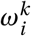, the mean number of offspring of a single individual of type *i* in a patch of habitat *k* (old or new). All adults are then removed.
iii. Mutation: the number of wild-type offspring mutating into the mutant type is drawn from a binomial distribution, with success probability *θ*, the mutation probability.
iv. Density regulation: if the number of offspring in a patch is higher than the local carrying capacity (*K_k_* for a patch of habitat-type *k* (old or new)), we sample *K_k_* individuals uniformly at random without replacement from the offspring population of the patch (hypergeometric sampling); all individuals have the same chance of survival at this step. Otherwise, the local population is left unchanged.

We consider that the mutant population has established if its total population size in patches of either the old- or the new-habitat type is greater than 60% the total carrying capacity of patches of that type ((0.6 × *K*_new_ × *M*(1 − *f*_old_)) for new-habitat patches, (0.6 × *K*_old_ × *Mf*_old_) for old-habitat patches).

Unless stated otherwise, the simulation results are averages of 10^5^ independent runs. All simulations are written in the C++ programming language and use the *Gnu Scientific Library*. The codes and data to generate the figures are deposited on Gitlab^1^.

## 3. Results

We proceed step-wise, as outlined above, towards the computation of the probability of evolutionary rescue. For each step, we first present a mathematical analysis, and then compare our results to the output of simulations that relax the assumptions made for mathematical purposes. First, we compute the *establishment probability* of a single mutant individual, depending on whether the mutation appeared in an old- or in a new-habitat patch, in an environment where the numbers of old- and new-habitat patches are fixed. Second, we derive an expression for the *probability of adaptation*, i.e. the probability for a mutation to appear in a given time interval and establish, again in a fixed environmental configuration. In this context, we also investigate the habitat of origin of the mutant lineages that eventually establish. Third, we study the time-varying scenario where patches, one after another, deteriorate, and we study the *probability of evolutionary rescue*. We again investigate the habitat of origin of the rescue mutant, and we compare the contributions to evolutionary rescue of standing genetic variation (i.e., mutations that are present before the environment starts deteriorating) and *de-novo* mutations (i.e., mutations that appear while the environment is deteriorating).

### 3.1. Establishment probability in a heterogeneous environment

In this first step, we consider that there is initially a single mutant individual in the population, located either in an old- or a new-habitat patch, and we compute the probability of establishment of the mutant population. In this step, we ignore further mutations and are only concerned with the fate of this single mutant lineage.

#### 3.1.1. Mathematical analysis

We denote by *φ_old_* (resp. *φ*_new_) the probability of establishment of this two-type branching process when the mutant is initially located in an old- (resp. new-) habitat patch. This probability is computed by considering all possible ways of going extinct: the initial individual having *j* successful offspring in a patch of type *k*, but all lineages descending from these *j* offspring eventually go extinct; then summing over *k* and *j*. Denoting by 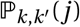 the probability that an individual in a *k*-habitat patch at the beginning of the generation has *j* successful offspring in a *k′*-habitat patch after density regulation, the following system of equations holds:

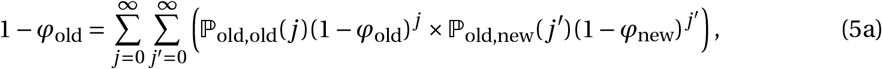

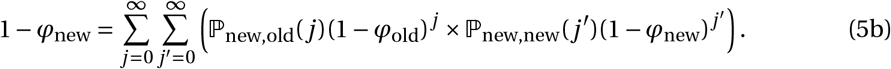

As mentioned previously, we assume for our mathematical analysis that the numbers of successful offspring per parent over the whole life cycle are Poisson distributed with means *λ_k,k′_* given in Eq. (4):

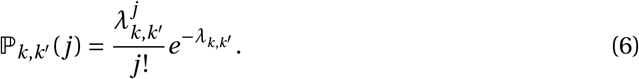

Inserting these expressions into system (5) and simplifying, we obtain

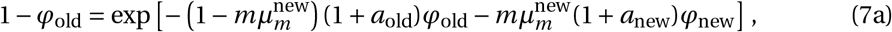

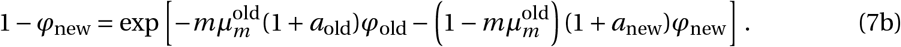

The establishment probabilities *φ_old_* and *φ_new_* are then given by the unique positive solution of system (7) (see Haccou et al., 2005, Chapters 5.3 and 5.6). This system of equations can be solved numerically. An analytical approximate solution is available in the case of weak selection and weak dispersal (i.e. *a*_old_, *a*_new_, *m* << 1); see for example Haccou et al. (2005, Theorem 5.6) for the general theory and Tomasini and Peischl (2018) for an application in a similar setting. The detailed derivation is presented in the SI, Section B. We find

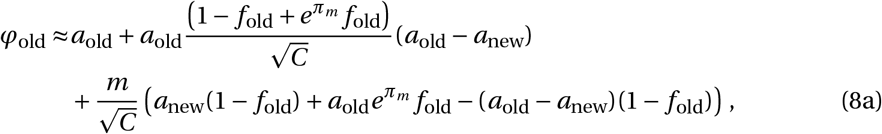

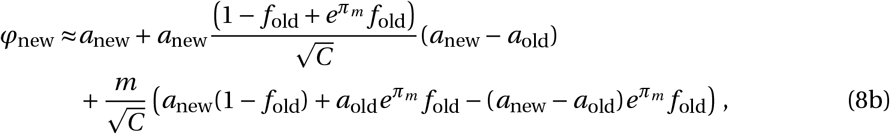

with

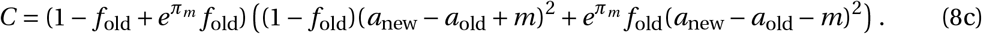

Recall that both *a*_old_ and *a*_new_, while considered constant in time, depend on the model’s parameters, and in particular on the dispersal probability *m*. The establishment probabilities *φ*_old_ and *φ*_new_ in (8) are therefore not affine functions of *m* (although they look so in Eqs. (8)).

If the emigration probability is zero (*m* = 0), the subpopulations in each habitat evolve in isolation from each other. The establishment probabilities in (8) become

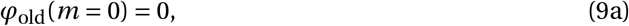

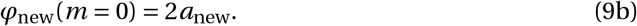

Eq. (9b) corresponds to Haldane’s classical result for the establishment probability of a slightly advantageous mutant (Haldane, 1927). The mutation being counter-selected in old-habitat patches, its probability of establishment is null (Eq. (9a)).

When the emigration probability is strictly positive (*m* > 0), in the case of unbiased dispersal (*π_w_ = π_m_* = 0) and for equal numbers of old- and new-habitat patches (*f*_old_ = 1/2), we recover the approximation found in Tomasini and Peischl (2018) (compare system (8) to their Eqs. (4) and (5)). Note that the approximation is independent of the actual number of patches (there are two patches in total in Tomasini and Peischl (2018)): the approximation only depends on the environmental configuration determined by the frequency of old-habitat patches *f*_old_.

#### 3.1.2. Comparison to simulations and qualitative behavior

Our mathematical analysis provided two kinds of results for the establishment probability: an implicit solution in Eq.(7), which we solve numerically, and an explicit but approximate solution in Eq.(8). In Fig. 2, we compare these solutions to simulation results for different values of the emigration rate *m*. We find good agreement with the numerical solution of Eq. (7) (solid lines). The approximation given in Eq. (8) (dashed lines) deviates slightly from the simulation results in regions where *m* is not small, i.e. when the assumptions made in the analytical derivation do not hold.

**Figure 2:**
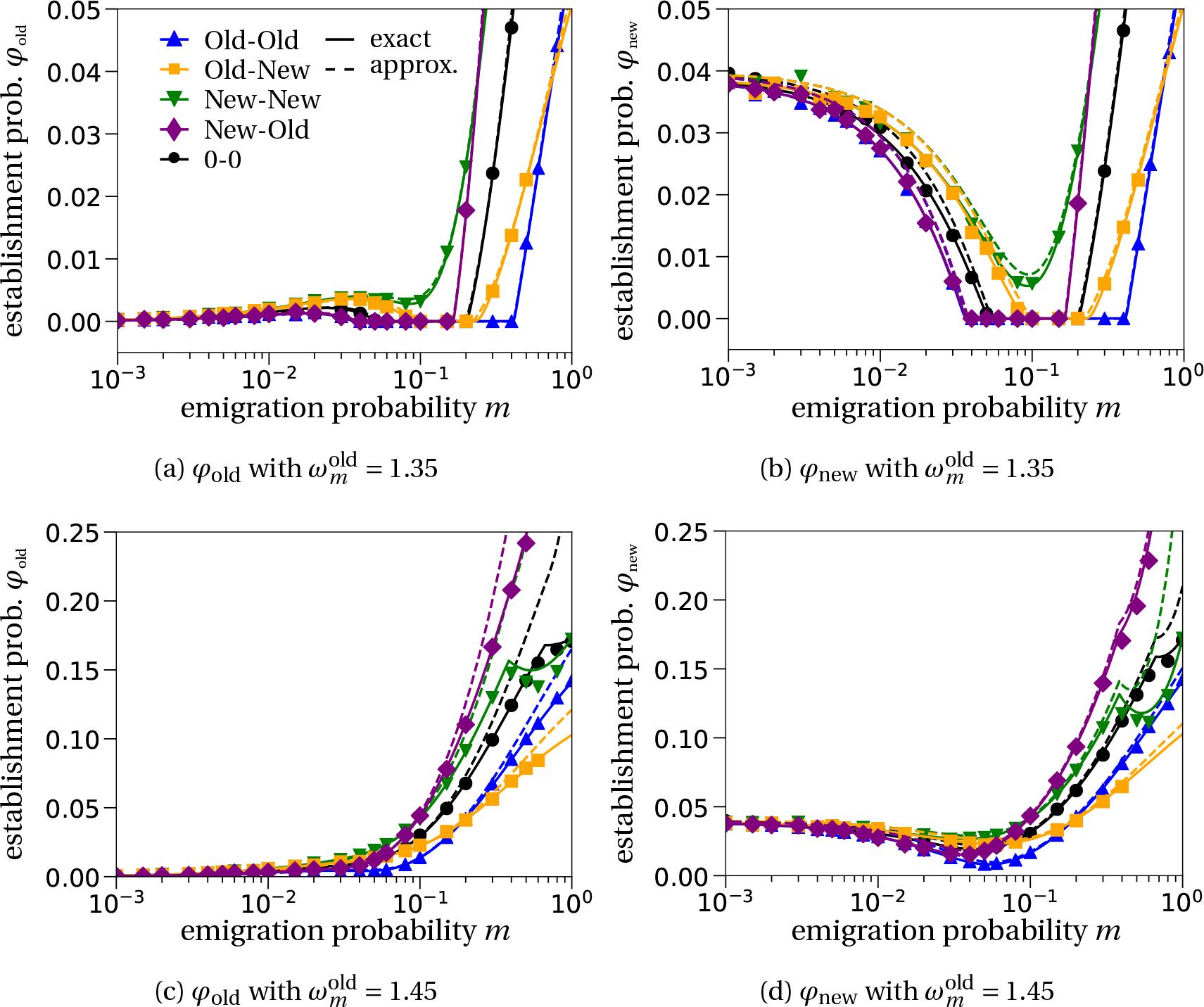
Establishment probability as a function of the emigration rate. Panels (a) and (c) show the establishment probabilities when the mutant arises in an old-habitat patch (*φ*_old_), and panels (b) and (d) the establishment probabilities when the mutant arises in a new-habitat patch (*φ*_new_), for mutant fecundity in old-habitat patches 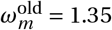 in (a), (b) and 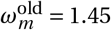 in (c), (d). Markers: simulations;full lines: numerical solution of Eq. (7); dashed lines, approximate solution shown in Eq. (8). The colors and marker shapes correspond to the different dispersal schemes, with the same parameters as in Fig.1. For a mutant emerging in old-habitat patches ((a),(c)), the establishment probability *φ*_old_ is 0 in the absence of dispersal (*m* = 0);it then increases with emigration *m*, which gives mutants a chance to settle in new-habitat patches where they are selectively favored;*φ*_old_ may still decrease at intermediate emigration probability (in (a)). For a mutant emerging in new-habitat patches ((b),(d)), the establishment probability *φ_new_* is approximately 2*a*_new_ = 0.04 in the absence of dispersal (*m* = 0). Increased dispersal is initially detrimental because mutants can land in old-habitat patches were they are selectively disfavored, but *φ*_new_ eventually increases with *m* thanks to competitive release in old-habitat patches. Large dispersal and a bias of the wild type towards the new habitat may inhibit the establishment of the mutant (New-New dispersal scheme in (c),(d)) because of gene swamping. For even larger *m*, competitive release becomes more important and the establishment probability re-increases.

We identify up to three different regions that define how the probability of establishment of a single mutant initially in an old-habitat patch (*φ*_old_) changes with the emigration probability *m* (Fig. 2a). This is in line with previous observations in the context of local adaptation (e.g. Kawecki, 1995; Tomasini and Peischl, 2018) and evolutionary rescue (Uecker et al., 2014). We define the regions as follows: (i) at low dispersal rates *m*, an initial increase of the establishment probability with *m*; (ii) a local maximum with a subsequent decrease of the establishment probability; (iii) at high dispersal rates *m*, an increase of the establishment probability with *m*.

In region (i), the beneficial effect of dispersal on the establishment probability *φ*_old_ is due to mutants dispersing from old-to new-habitat patches where they are fitter than the wild type. While this effect is still present in region (ii), the establishment probability *φ*_old_ now decreases with dispersal because the offspring of individuals that dispersed to a new-habitat patch can disperse back into old-habitat patches. More precisely, the expected per capita number of successful offspring in the new habitat of an adult in a new-habitat patch is 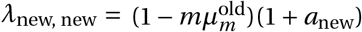. This product can, for large emigration probabilities *m*, be smaller than 1, i.e. a mutant in a new-habitat patch has on average less than one offspring in a new-habitat patch. This is detrimental to the mutant because it means that mutants do not efficiently reproduce in the habitat where they are fitter. Finally, in region (iii) at high dispersal, so many wild-type individuals leave old-habitat patches that competitive pressure in old-habitat patches is substantially decreased. The post-dispersal size of the wild-type population 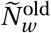 is then low enough that the local growth rate of the mutant in these patches, *a*_old_ (first term in Eq. (8)), becomes positive (Fig. S3). This effect is called ‘relaxed competition’ (Uecker et al., 2014). The onset of this effect, in terms of the emigration probability *m*, is strongly dependent on the difference in fecundity of the mutant and the wild type in the old habitat. The smaller the difference in fecundity is, the ‘earlier’ (i.e. for smaller emigration rates *m*) relaxed competition becomes relevant (compare panels 2a to 2c), to a point that region (ii) vanishes (panel 2c) and there is no decrease of *φ*_old_ with *m* any longer. In contrast, for lower mutant fecundity values 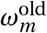, region (iii) might vanish (see Fig. S4 in SI), because the mutant’s fecundity in old-habitat patches is too low compared to the wild type’s, so the mutant does not benefit from relaxed competition in old-habitat patches.

The qualitative behavior of the establishment probability of a mutant emerging in the new habitat, *φ*_new_, can be interpreted in a similar way (panels 2b, 2d). The establishment probability *φ*_new_ decreases with the emigration probability *m* at low *m*, because the mutant appeared in a new-habitat patch, where it fares better than the wild type, so there is no initial benefit due to dispersal. When the emigration probability is higher, however, mutants can back emigrate to new-habitat patches, while those that land in old-habitat patches can enjoy relaxed competition when *m* is high. This is why *φ*_new_ increases with *m* at higher *m*.

An additional effect can take place at high dispersal and reduce mutant establishment probabilities, in particular when the wild type is biased toward new-habitat patches (see for instance the New-New scheme in Figs. 2c,d). The high dispersal of wild-type individuals lets the local population in new-habitat patches exceed the carrying capacity *K*_new_, inhibiting the establishment of a locally better adapted type (mutant). Note that the lower carrying capacity in new-habitat patches than in old-habitat patches (*K*_new_ < *K*_old_) creates a favorable setting to this effect, also referred to as gene swamping (Nagylaki, 1978; Lenormand, 2002). Further increasing the emigration probability *m* results in relaxed competition in the old habitat, which explains the re-increase of the New-New dispersal scheme.

#### 3.1.3. Comparison of dispersal schemes

We now compare the establishment probabilities across the different dispersal schemes. Mostly, a general bias towards the new habitat (New-New in Fig. 2) enhances mutant establishment compared to the unbiased dispersal scheme (0-0). This can be attributed to two reasons. First, the mutant is more likely to disperse to the new habitat where it outcompetes the wild type. Second, competition in old-habitat patches is relaxed starting at low emigration probabilities *m* because the wild type preferentially disperses to new-habitat patches. Conversely, a bias towards the old habitat (Old-Old) lowers mutant establishment probabilities compared to the unbiased dispersal scheme.

The rankings of the type-dependent dispersal schemes (Old-New and New-Old) compared to the unbiased scheme (0-0) depend on the amount of dispersal (compare the orange, purple and black curves in Fig. 2). As explained above, at low dispersal probabilities *m*, the prevalent force is the dispersal of mutants to new-habitat patches. The establishment probability of the mutant is therefore higher for the Old-New scheme, where mutants preferentially disperse to new-habitat patches, as compared to random dispersal, while the opposite is true for the New-Old scheme. At high dispersal probabilities *m* however, an important force is competitive release in old-habitat patches. The establishment probability of the mutant is therefore higher in the scheme where wild-type individuals preferentially disperse out of old-habitat patches, releasing competition there (New-Old scheme).

### 3.2 Probability of adaptation in a heterogeneous environment

We now study the probability of adaptation when mutations occur recurrently. As in the previous section, we consider a heterogeneous environment with a fixed number of old- and new-habitat patches. This is effectively a source-sink system (Holt, 1985; Pulliam, 1988), where old- and new-habitat patches correspond to sources and sinks for the wild type, respectively. In the previous section, we initialized the system with one mutant in either an old- or a new-habitat patch and computed the establishment probability. Now, we let mutants appear randomly within a certain time frame. The last time point at which a mutation can occur is denoted by *t*_fin_.

In this setting, the probability of adaptation *P*_adapt_ is approximated by

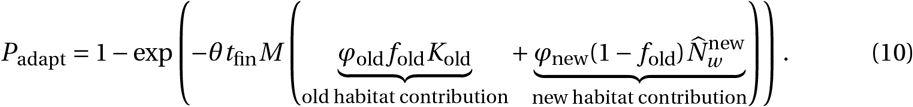

This is one minus the probability that no mutant establishes within the [0, *t*_fin_] time interval, given by the probability of zero successes of a Poisson distribution. The rate of this Poisson distribution is the expected number of successfully emerging mutant lineages until time *t*_fin_.Mutants arise with probability *θ*; *M*_old_*K*_old_ is the total number of wild-type individuals in old-habitat patches, 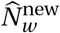 the total number of wild-type individuals in new-habitat patches. A mutant arising in a *k*-habitat patch has a probability *φ_k_* of establishing in the population; we assume that mutants establish independently of one another. Assuming a Poisson distribution for the number of successful mutant lineages is an approximation of a Binomial distribution with large sample size (the wild-type population size) and small success probability (the establishment probabilities *φ_k_* times the mutation probability *θ*). Note also that for *t*_fin_ tending to infinity, there will almost surely be a successful mutant, so that *P*_adapt_ = 1.

The probability of adaptation *P*_adapt_ calculated with Eq. (10) is compared to simulation results in Fig. 3. In spite of our approximations, the fit to simulation results is still very good. As *P*_adapt_ includes the probabilities of establishment *φ_k_*, here again, the shapes of the curves as function of the emigration probability *m* depend on the fecundity of the mutant in old-habitat patches, 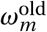 (Figs. 3a, 3c). Likewise, the rankings of the different dispersal schemes are comparable to the ones observed for the establishment probability.

**Figure 3:**
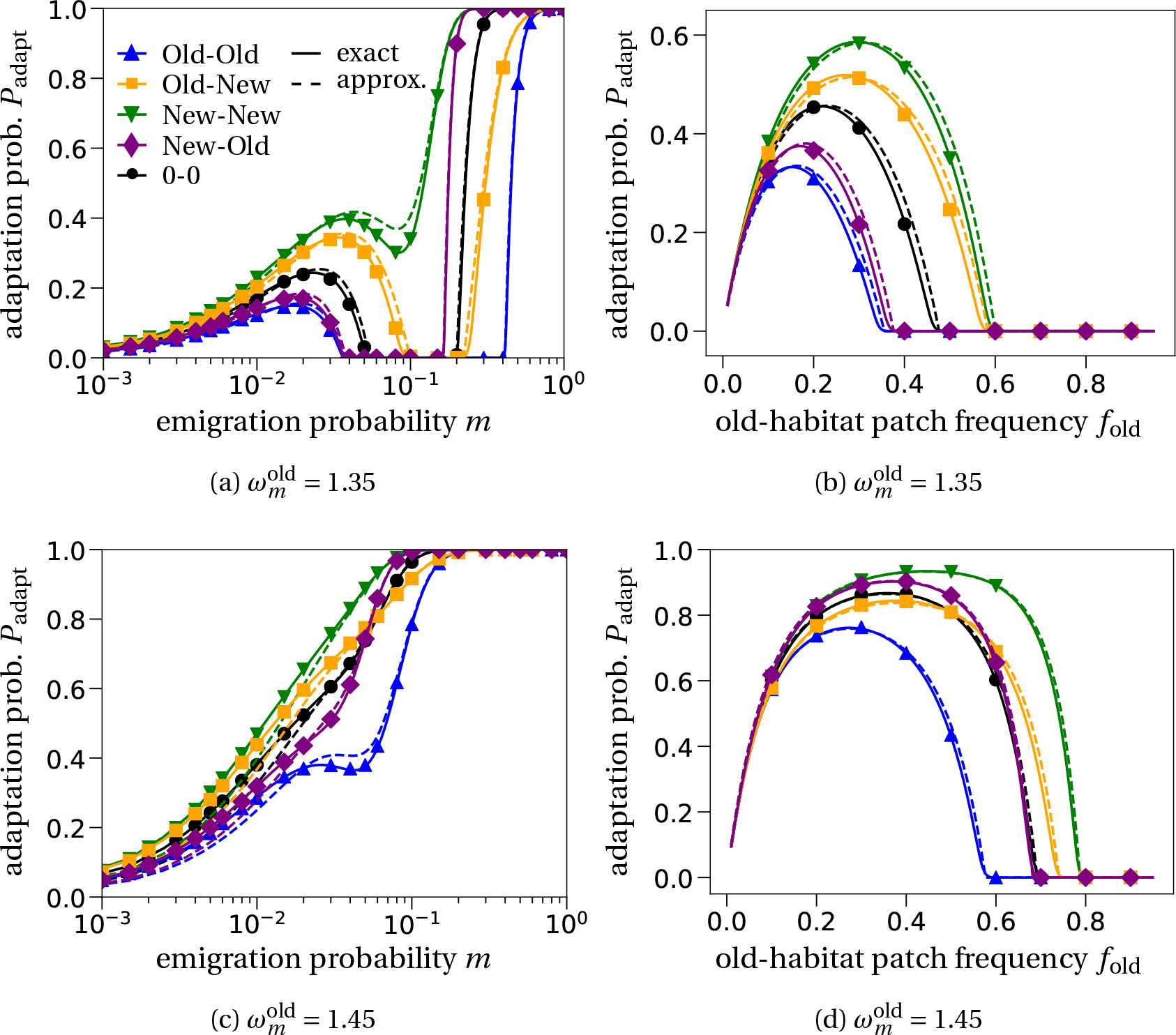
Probability of adaptation in a heterogeneous environment. In (a) and (c), we vary the emigration rate *m* and observe a similar qualitative behavior as for the establishment probability *φ*_k_ in Fig. 2. In (b) and (d), we vary the frequency of old-habitat patches. The maximum is the result of two counteracting processes. The higher the number of old-habitat patches (the greater *f*_old_), the larger the wild-type population. As a consequence, more mutants appear in the studied time-frame. In contrast, the less old-patch habitats there are in the environment (the lower *f*_old_), the higher the probability of successful establishment of a mutant population. The curves are given by Eq. (10); ‘approx.’ curves use the approximate solution for *φ_k_* given in Eq. (8), while ‘exact’ curves are obtained by numerically solving Eq. (7) for *φ_k_*. In all panels, the mutation probability is *u* = 1/(*MK*_new_) and the final time for a mutant to appear is *t*_fin_ = 100.

Panels 3b and 3d show the probability of adaptation as a function of the frequency of old-habitat patches *f*_old_. The maximum of *P*_adapt_ at intermediate *f*_old_ is the result of two antagonistic effects: (1) the likelihood that a mutation appears increases with the number of wild-type individuals present in the system, which is highest for high frequencies of old-habitat patches *f*_old_, and (2) the probability of establishment of a mutant decreases with the number of old-habitat patches.

The different dispersal schemes alter both effects. The probability of adaptation is highest when there is a general bias towards the new habitat (New-New), due to a combination of high establishment probabilities (Fig. 2) and high local population sizes thanks to the bias (Fig. S1). Conversely, a general preference for old-habitat patches (Old-Old) yields lower probabilities of adaptation.

#### 3.2.1 Habitat of origin of the adaptive mutation

We now ask in which habitat mutations leading to successful establishment appear. To do so, we distinguish in the simulations between mutants that appear in old-habitat patches and mutants that appear in new-habitat patches, and identify the habitat of origin of the mutation by considering the composition of the mutant population after establishment. We however do not distinguish between separate mutations that appear in the same type of habitat, meaning that we cannot rule out the presence of multiple lineages if they all appeared in the same type of patch: there may be soft selective sweeps (see Hermisson and Pennings, 2017, for a review). With this implementation, it is only if the established mutant population contains mutants that appeared in both old- and new-habitat patches can we be sure that multiple lineages contributed.

Analytically, we approximate the probability to observe a mutant population that can be traced back to mutants from old-habitat patches only by

P(successful adaptation from old habitat)(1-P(successful adaptation from new habitat))

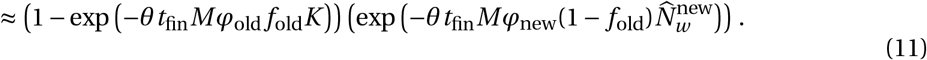

The corresponding probabilities for the other two scenarios can be computed analogously. The approximation uses our key assumption that different mutant individuals and their offspring do not affect each others’ dynamics (branching process). In the simulations, we label a run as having established lineages originated from different habitat types (“both” in Fig. 4) if these lineages are still alive alter 1,000 generations (rather than counting the number of lineages right after the mutant population size has crossed the establishment threshold). This lowers the likelihood that we count false-positives where a mutant in one of the habitats has just arisen right before the mutant population exceeds the establishment threshold. Simulations where established lineages arose exclusively in old- or new-habitat patches are labeled “old-habitat patch” and “new-habitat patch”, respectively.

**Figure 4:**
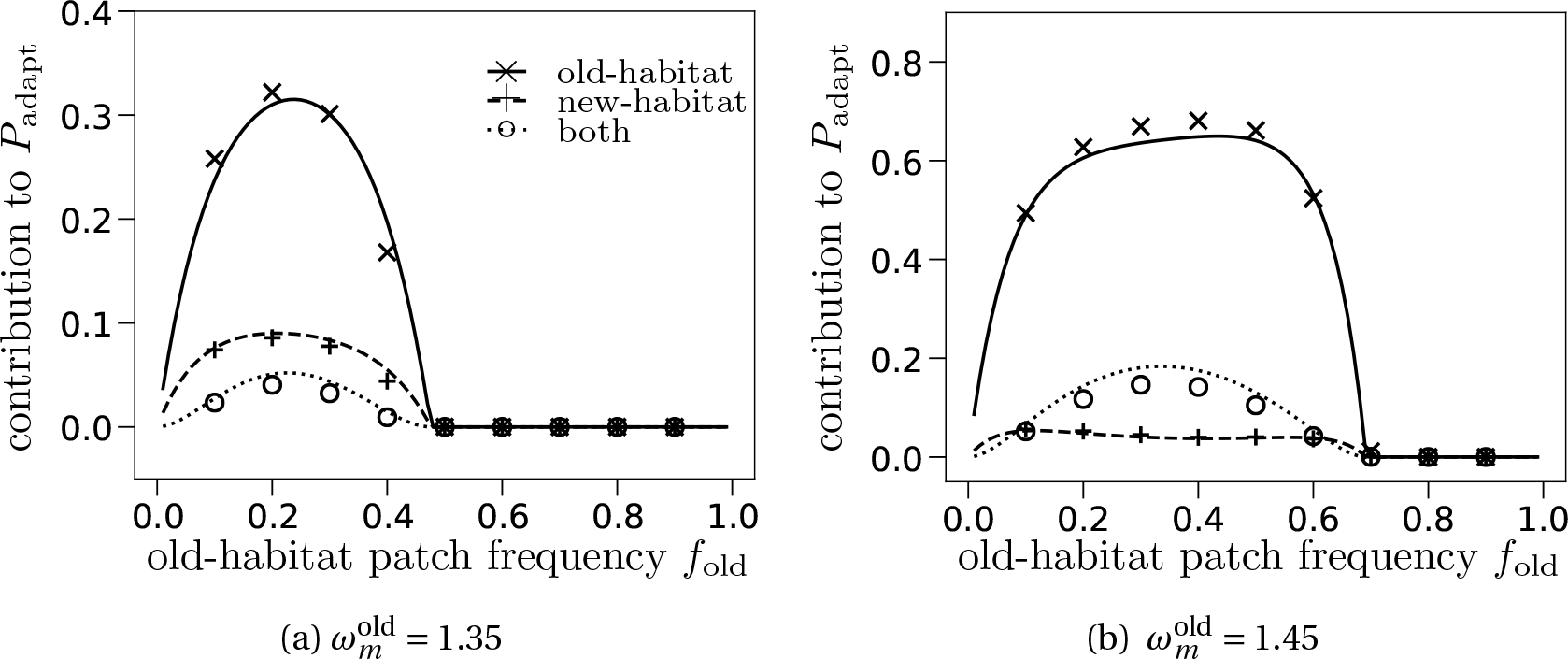
Origin of the adaptive mutant, depending on mutant fecundity in old-habitat patches 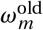 (recall that 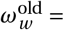 1.5). The points correspond to simulations, where mutants arising in old-vs. new-habitat patches are differently labeled, and where we consider the composition of the mutation population at the end of the simulation. The labels “old-habitat” and “new-habitat” correspond to established mutant lineages from exclusively that habitat type, “both” refers to mutant populations that can be traced back to both habitat types. The curves are given by Eq. (11) (or the adjusted versions of it) under the unbiased dispersal scheme (*π_w_ = π_m_* = 0). Note the different scaling on the y-axes. Mutants that establish predominantly appeared in old-habitat patches: the lower establishment probability for mutants emerging in old-habitat patches is compensated by the larger number of mutants appearing in these patches, due to a higher local population size.

We compare our calculations to simulation results in Fig. 4, varying the frequency of old-habitat patches *f*_old_. Most successful mutations arise in old-habitat patches, with a much smaller contribution to the probability of adaptation for lower number of mutant offspring in old habitats (Fig. 4a) than for larger numbers of offspring (Fig. 4b). The contributions of old-vs. new-habitat patches depend on the product 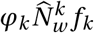, which we decompose in Fig. S5. Even though the establishment probability from old-habitat patches is lower (*φ*_old_ < *φ*_new_), the total population size of the wild type in old-habitat patches is larger than that in new-habitat patches, so that more mutants appear in old-habitat patches than in new-habitat patches, which compensates their lower establishment probability.

### 3.3. Evolutionary rescue

Finally, we assume that patches deteriorate one after another at regular time intervals *τ*, until all patches have switched to the new habitat. If the wild-type population fails to generate a successful mutant, the population will inevitably go extinct, because the entire environment will consist of new-habitat patches, and because a wild-type population is assumed not to be viable there. We first focus on evolutionary rescue due to *de-novo* mutations. Because the configuration of the environment changes overtime, we denote by *f*_old_(*i*) = (*M* – *i*)/*M* the proportion of old-habitat patches after the *i*^th^ deterioration event. We also explicitly write the dependency of the establishment probabilities on the proportion of old-habitat patches, *φ_k_*(*f*_old_(*i*)) – this is only a notation change, the formulas presented before still apply (Eqs.(7) and (8)). We approximate the probability of evolutionary rescue, denoted by *P*_rescue_, as

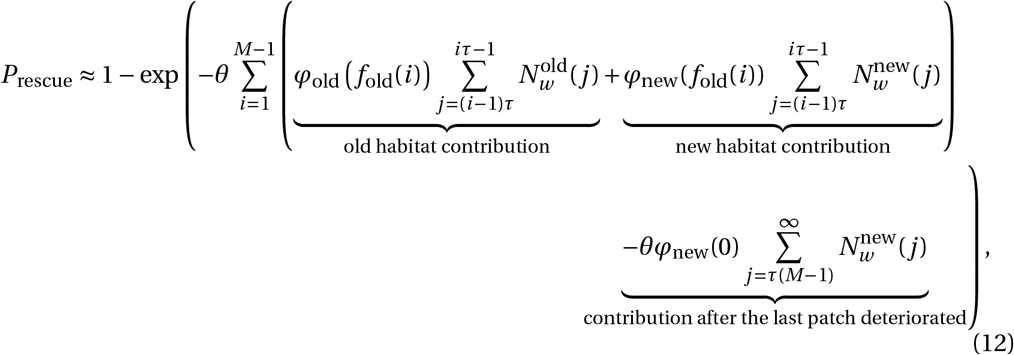

where 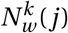 denotes the overall number of wild-type individuals living in habitat *k* (old or new) in generation *j* (see SI, Section A.4, Eq. (A.8)). The interpretation of this equation is the same as for the probability of adaptation in Eq. (10). The only difference is that we now need to account for a changing environment. In the formula, these changes are accounted for by the sums that iterate through the (discrete) time steps and by the time dependence of the corresponding quantities. We further note that we follow the expected value of the wild-type population size deterministically over time 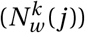, instead of assuming it to be already in its steady state as in Eq. (10). The establishment probabilities *φ_k_*(*f*_old_(*i*)) are however still computed using stationary population sizes, calculated at each time step using Eqs. (7) and (8) (i.e. they are considered as piecewise constant).

Comparison to simulated data indicates that the approximation in Eq. (12) correctly predicts the ranking of dispersal schemes; the actual fit to data is however less good than for the previous steps of our analysis. This discrepancy can be explained: our analysis assumes that for a mutant born in a certain patch configuration, say with *i* old-habitat patches, the environment does not change anymore. That is, a mutant born in a *k*-habitat patch in this environment contributes *φk(i/M*) to the probability of evolutionary rescue despite further patches deteriorating—while having more new-habitat patches increases the probability of establishment, see for example Fig. S5. Thus, the probability of establishment is underestimated. This is especially true for mutants that emerge just before a deterioration event. Additionally, *φ_k_(i/M*) assumes stationary wild-type population sizes and therefore does not reflect the decreasing wild-type population size right after the deterioration of a patch. A time-dependent establishment probability could account for these effects but unfortunately is not amenable to approximations in our framework. Uecker et al. (2014) were able to find a time-dependent solution by focusing on specific scenarios: situations with either full mixing of the global population (*m* = 1) or a sterile mutant in old-habitat patches 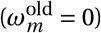. In these extreme cases, the branching process becomes one dimensional, and an analytical, time-dependent solution can be obtained, which is not the case with a two-dimensional branching process like ours.

The ranking of the different dispersal schemes is overall conserved from the previous steps of our analysis (Fig. 3). Differences between the dispersal schemes are more marked when the fecundity of the mutant in old-habitat patches is lower (Fig. 5c,d), including when the mutant cannot reproduce at all in old-habitat patches 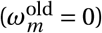. It is comparatively better for rescue that the mutant preferentially disperses into new-habitat patches, where it is relatively fitter, and for the wild type to also preferentially disperse into new-habitat patches, thereby releasing competition in old-habitat patches.

**Figure 5:**
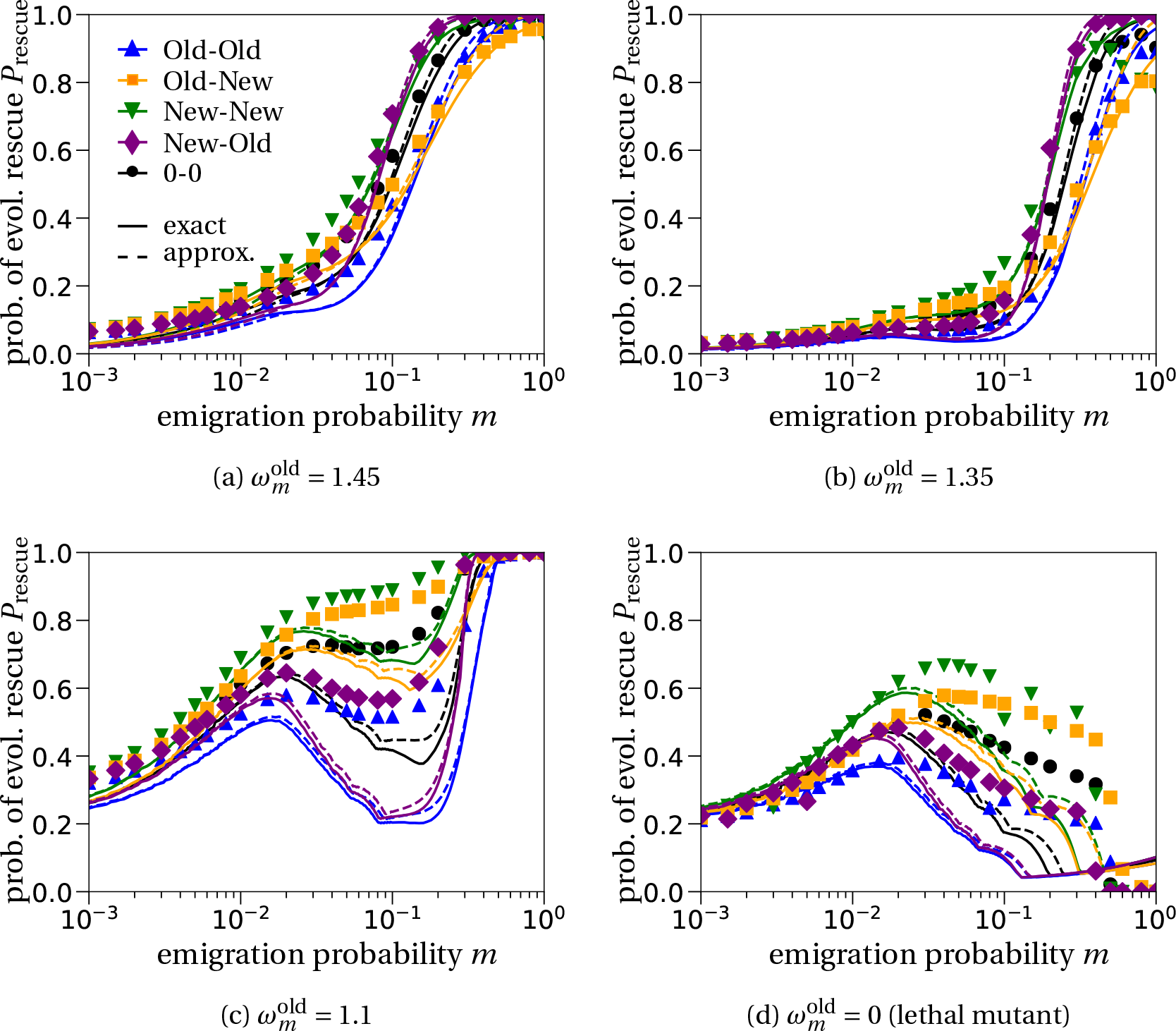
The probability of evolutionary rescue compared to simulation results. Our predictions, computed with Eq. (12), match the qualitative behavior of the simulated data for the probability of evolutionary rescue. All rankings of the dispersal schemes align well. Quantitatively though, we find that our predictions tend to underestimate the simulated data. In (a,b) the mutation probability is set to *θ* = 1/(25*MK*_new_), in (c,d) it is *θ* = 1/(*MK*_new_). For the establishment probabilities *φ_k_* in Eq. (12), the solid lines show the exact solution of Eq. (7) and the dashed lines show the approximated (‘approx’) solution from Eq. (8).

When mutant fecundity in old-habitat patches is comparatively low (Fig. 5c,d), the probability of evolutionary rescue often reaches a local (or global) maximum at intermediate emigration probabilities. This finding extends previous results (Uecker et al., 2014; Tomasini and Peischl, 2020) to arbitrary dispersal schemes affecting the immigration process.

#### 3.3.1. Habitat of origin of the rescue mutant and standing genetic variation

Similar to what we found for the probability of adaptation, rescue mutants mainly originate from old-habitat patches (Fig. 6a). Mutations are more likely to appear in the more populated patches (old habitat). However, a low mutant fecundity in old-habitat patches 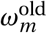 decreases the chance of establishment of these mutants that appear in old-habitat patches (compare black and yellow symbols in Fig. 6a). Here again, we cannot rule out that multiple mutant lineages having appeared in the same habitat type established. Only when mutants from both habitat types are present can we be sure that at least two lineages contributed to establishment (i.e., there was a soft sweep). In our parameter set, the probability of rescue with a mix of origins was very low in our simulations (circles in Fig. 6a). Note that our choice of a small mutation rate corresponds to a hard selective sweep regime (*θK*_old_*M* = 0.08 < 1) (Wilson et al., 2017a; Hermisson and Pennings, 2017).

**Figure 6:**
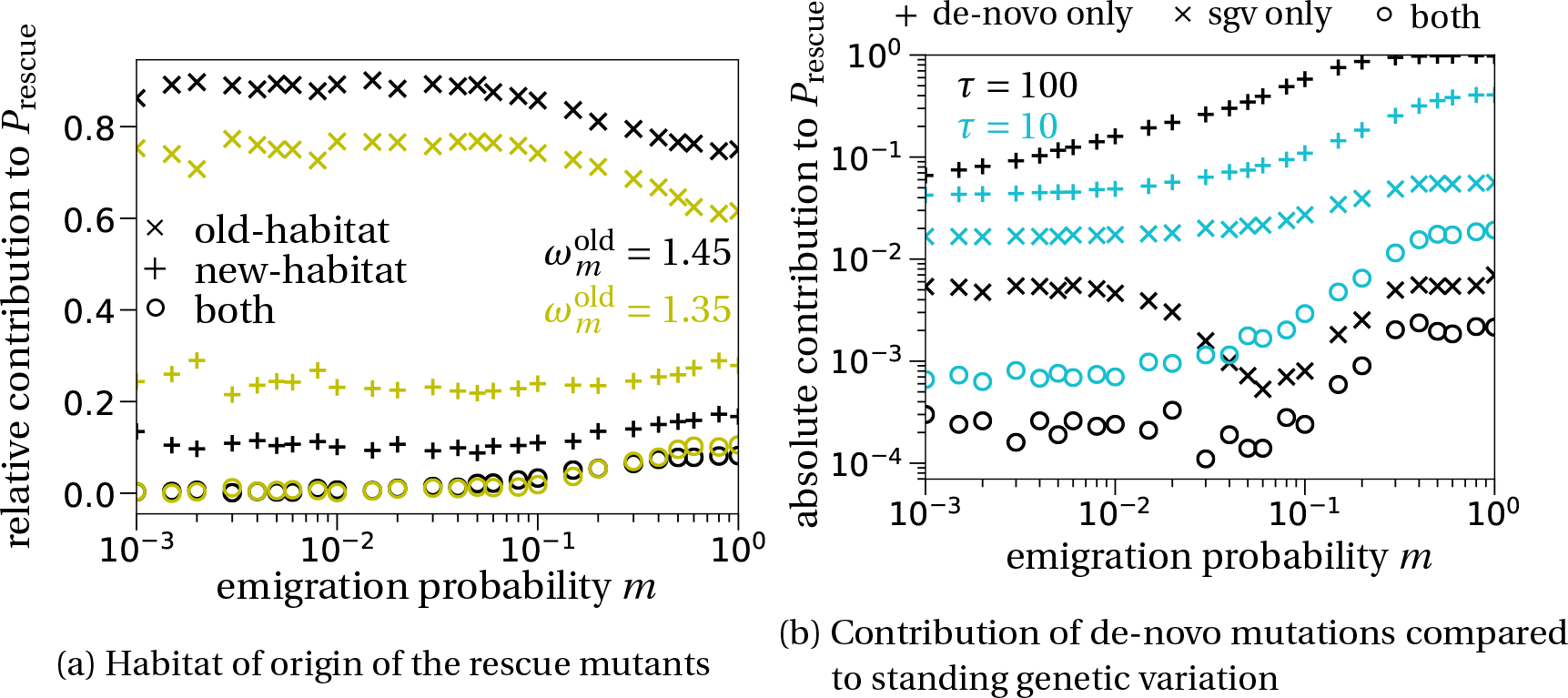
Habitat of origin of the rescue mutation and the impact of standing genetic variation. (a) We compare the origin of successful mutations for different mutant fecundity in the old-habitat patches (black vs. yellow). Comparatively more established mutant originated from new-habitat patches when the mutant fecundity in old-habitat patches 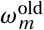 is lower (compare black and yellow + markers). Dispersal is unbiased (*π_m_ = π_w_* = 0). (b) The larger the time interval *τ* between two degradation events, the smaller is the influence of standing genetic variation on the probability of evolutionary rescue (compare black and blue × markers). For large emigration probabilities, the probability of evolutionary rescue due to de-novo mutants increases (Fig. 5). This is also true for rescue events due to standing genetic variation (see blue × markers). The simulations are done by letting the system evolve for 1,000 generations before the first deterioration event happens. Parameters: *π_m_ = π_w_* = 0 in both scenarios and 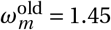. Note the log-scale on the y-axis.

So far, we have considered settings where evolutionary rescue is exclusively due to de-novo mutations. To explore the role of standing genetic variation (sgv), we ran simulations where we let the system evolve for 1,000 generations before the first degradation event happened – we were not able to find a theoretical prediction for the contribution of standard genetic variation. Mutants that appeared before the first degradation event, i.e. at times *t* < 0 when *f*_old_ = 1, were labeled ‘sgv-mutants’. Mutant appearing after *t* = 0 are labeled ‘de-novo’ mutants. Fig. 6b shows the contributions of de-novo mutations and of standing genetic variation to the probability of evolutionary rescue. Rescue events involving mutants from standing genetic variation are much rarer than rescue events from de-novo mutants (note the log scale in Fig. 6b). The number of rescue events due to standing genetic variation decreases when the interval between two degradation events, *τ*, increases (compare blue and black × markers in Fig. 6b; see also Fig. S6 in Section E in the SI for more details). This is because mutants that were present at time *t* = 0 (sgv-mutants) needed to survive for sufficiently many patch deterioration events before their growth rate turned positive, giving them a chance to establish. The longer this time (higher *τ*), the less likely their establishment. As already shown before, the probability of evolutionary rescue by de-novo mutations increases with emigration probability *m* when *m* is large (Fig. 5). This is also the case for ‘sgv-mutants’ (Fig. 6b), for the same reasons (relaxed competition in old-habitat patches when *m* is large).

## 4. Discussion

We have studied the probabilities of establishment, adaptation and evolutionary rescue under four non-uniform dispersal schemes and compared them to unbiased dispersal. In line with previous results, we find that the probabilities of establishment, adaptation and evolutionary rescue can display up to three different phases when varying the dispersal rate *m*. The dispersal schemes affect population dynamics and consequently the parameter regions corresponding to the three phases.

### 4.1 Dispersal and adaptation

Theoretical studies that investigated the effects of spatial subdivision on the adaptation of a population in a heterogeneous environment can be classified into two types, depending on how they treat demography. One type of models, classically analyzed in a population genetic framework, assumes constant population sizes in all patches, independent of their local habitat type and of dispersal strength (a feature which we later call implicit demography). Results obtained in this framework show one notable difference when compared to our model with demography: in these models, the probability of successful establishment of a rare mutant favored in some part of the environment decreases at larger dispersal rates (e.g. Nagylaki, 1978; Bürger and Akerman, 2011). This gene swamping effect is due to the dispersal of non-adapted individuals into the habitat type where the rare mutant is beneficial, decreasing the local frequency of the mutant (Lenormand, 2002; Tomasini and Peischl, 2018).

The second type of models explicitly takes into account demographic effects due to dispersal, often in the context of source-sink systems (Holt, 1985; Pulliam, 1988). Here, the effect of dispersal on adaptation depends on the growth rate differences of the mutant and the wild type in the two habitats (Kawecki, 2000), which we also observe. When the mutant is just slightly less fit than the wild type in the old habitat (Fig. 2c), the probability of adaptation monotonically increases with dispersal. When the mutant’s fecundity is lower, establishment probabilities reach a local maximum at intermediate dispersal rates, and increase again at large dispersal rates thanks to relaxed competition (Fig. 2a(a)). When the fecundity of the mutant is even smaller, the local maximum remains but relaxed competition no longer occurs (cf. Fig. S4).

To compare the effects of explicit vs. implicit demographic dynamics, we provide in the SI a non-demographic version of our model (Section F and Fig. S7), i.e. where all patches are at carrying capacity. Relaxed competition can also happen in models with implicit demography, but only if the dispersal preference is type-dependent (New-Old or Old-New; Fig. S7). This is because, for large emigration probabilities, type-dependent dispersal preferences cause a quasi-separation of the mutant and the wild type into different patch types, so they are less directly competing.

### 4.2. Standing genetic variation and evolutionary rescue

We also studied the contribution of standing genetic variation to evolutionary rescue. This contribution increases with the speed of environmental change (i.e., with smaller intervals between degradations *τ*), Figs. 6b and S6. This observation has also been made in a quantitative genetics setting where the adaptive trait is continuous (and not discrete as in our model) (Matuszewski et al., 2015). Experimental results with *Caenorhabditis elegans* also indicate that the impact of standing genetic variation is small under slow environmental change (Guzella et al., 2018).

### 4.3. The effect of biased dispersal patterns on adaptation and evolutionary rescue

The importance of considering dispersal schemes other than unbiased dispersal has been highlighted in several papers (Edelaar et al., 2008; Clobert et al., 2009; Edelaar and Bolnick, 2012). This has led to a number of simulation studies exploring the effect of various dispersal schemes on (local) adaptation (e.g. Vuilleumier et al., 2010; Holt and Barfield, 2015; Mortier et al., 2018; Pellerin et al., 2018). These cited studies examined the effect of matching habitat choice on adaptation in a heterogeneous environment and observed that it increases the probability of adaptation compared to unbiased dispersal.

We identified two types of effects of the different dispersal schemes. First, by changing population densities in both habitat types, the dispersal schemes alter the growth rate of the mutant in both patch types (Fig. S3). This is the primary reason for the ranking of the dispersal schemes, with a general immigration bias into new habitats (New-New) promoting evolutionary rescue the most and a general immigration bias into old habitats (Old-Old) promoting it the least. Second, the different dispersal schemes affect the number of mutations arising in either habitat type. This has a minor effect on the probability of evolutionary rescue for the explored parameter range but is relevant for the origin of the successful mutant lineage (see also Fig. S8). As the genetic background may vary across patches, the origin of a successful mutation will also affect which mutations will hitchhike with it. Similarly, with polygenic rescue or under recombination (e.g. Schiffers et al., 2013; Uecker and Hermisson, 2016), the origin of a mutant is likely to affect its success, as is the case in our model.

### 4.4. Generality of our theoretical analysis and future directions

Our mathematical results rely on the simplifications that the wild type population does not change over time and that the mutant is rare enough that mutants live independently of each other, and do not affect wild-type individuals. This allows us to summarize mutant population dynamics with the *λ* terms presented in Eq.(4). Furthermore, for our approximation in Eq. (8) to generate accurate predictions, it is essential that growth rate differences between the wild type and the mutant are weak and dispersal is low – these conditions are however not needed when we numerically solve system (7) (see also Section H in the SI where we relax the condition of small mutant fecundity in the new habitat).

Our approach can account for various dispersal schemes and local type-dependent population dynamics, i.e. different reproduction and competitive parameters for the wild type and the mutant. However, it cannot account for type-dependent carrying capacities, explicit spatial structure or rapidly changing environments.

Lastly, our model can readily be extended and include a cost of dispersal or a different life cycle. In particular, the variation of the life cycle could yield distinct results regarding adaptation (Débarre and Gandon, 2011; Holt and Barfield, 2015) and, more generally, in the context of the evolution of dispersal (Massol and Débarre, 2015).

### 4.5. Conclusion

In conclusion, we studied the effect of dispersal and habitat choice on the probability of establishment, adaptation and evolutionary rescue of a mutant under divergent selection in a subdivided population. We recover previous results on adaptation and, using results from the theory of multi-type branching processes, provide a general framework for studying evolutionary dynamics of a subdivided population in heterogeneous environments in discrete time. This unifying approach allows us to identify the forces responsible for the different predictions obtained in the population genetics literature and under source-sink dynamics. We find that including population demography significantly alters the results for high dispersal rates. For constant population sizes and type-independent dispersal schemes, high dispersal rates have a negative effect on establishment, while with explicit demography the effect is largely positive. The latter is a result of relaxed competition in old-habitat patches. Most importantly, we extend the existing literature by comparing different dispersal schemes and studying their effects on adaptation and evolutionary rescue. We find that a general dispersal bias towards degraded patches, New-New dispersal scheme, results in the highest probability of adaptation and evolutionary rescue. The lowest values are obtained for the Old-Old dispersal scheme. These results show that non-uniform dispersal patterns can have a strong influence on population survival and adaptation in a heterogeneous environment.

## Acknowledgements

PC and FD received funding from the Agence Nationale de la Recherche (ANR-14-ACHN-0003 to FD). HU appreciates funding from the Max Planck Society. We are grateful to the INRAE MIGALE Bioinformatics Facility (MIGALE, INRAE, 2018. Migale Bioinformatics Facility, doi: 10.15454/1.5572390655343293E12) for providing computational resources. We thank Jérôme Mathieu for highlighting the connection of the ‘New-Old dispersal scheme’ to the ecological trap literature, and Staffan Jacob and Pim Edelaar for fruitful discussion concerning the biological motivation of the dispersal schemes. We thank two anonymous reviewers and Associate Editor David Vasseur for thorough comments that helped us improve our manuscript.

## Appendix

## A. Deriving the model dynamics

In this section, we provide the mathematical details of the model that is verbally described in the main text. We start by deriving the population dynamics when only the wild type is present. This will allow us to compute the local growth rate of a rare mutant.

Before we go into the details of the computation, we recall the form of the dispersal rates. A dispersing wild-type individual immigrates to a new-habitat patch with probability

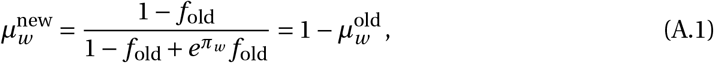

where *f*_old_ is the frequency of old-habitat patches and *π_w_* is the wild-type bias towards old-habitat patches. The complement, 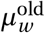, is the probability that the dispersing wild-type individual instead immigrates into an old-habitat patch.

All the subsequent computations can be checked with a symbolic programming language (e.g. *Mathematica*). A *Mathematica* notebook is deposited on Gitlab^2^.

### A.1. Stationary wild-type population sizes

We denote by 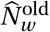 and 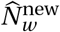 the deterministic stationary population sizes of the wild type in old- and new-habitat patches. We assume that the population is always at carrying capacity in old-habitat patches, so that 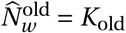. We compute 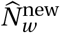 recursively. It is given by the solution of the following equation:

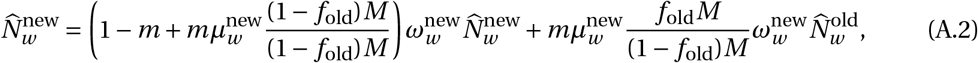

where the first term on the right-hand side corresponds to individuals born in a new-habitat patch and staying in it or migrating and landing in a new-habitat patch, and the second term corresponds to individuals born in an old-habitat patch and migrating to a new-habitat patch. Simplifying, using Eq. (A.1) to replace 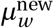 and replacing 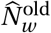 by 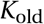, we obtain

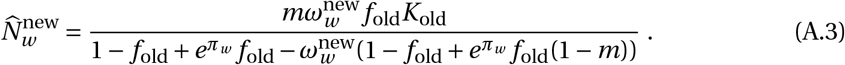

This value cannot be larger than *K*_new_, the carrying capacity of new-habitat patches. So,

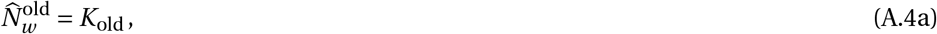

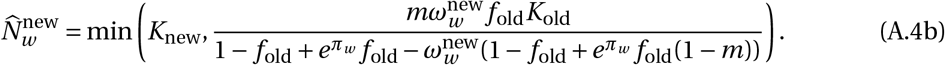

The shape of 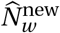 as function of *m*, with the parameters used in the main text, is given in Fig. S1.

**Figure S1:**
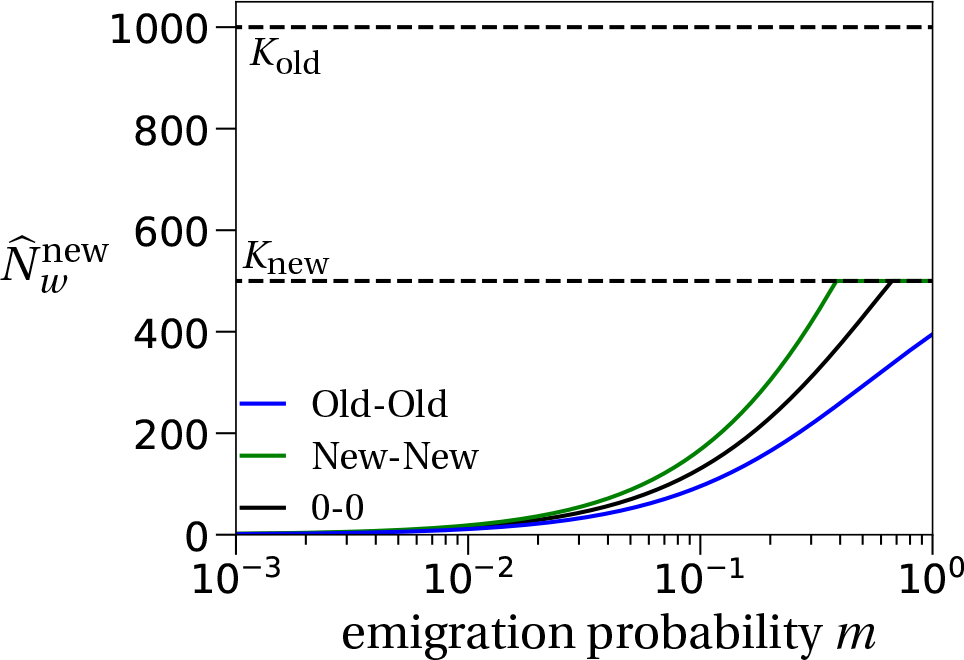
Stationary wild-type populations. The stationary wild-type populations as a function of the emigration probability *m* only depend on the wild-type bias on immigration. Therefore, we do not plot the curves for the asymmetric dispersal schemes, Old-New and New-Old. The stationary wild-type population size in old habitats, 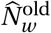 is constant for all values of the emigration probability *m*. The parameters are given in Table 1 with 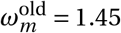.

### A.2. Wild-type population sizes after dispersal

We denote by 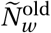 and 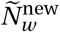 the numbers of wild-type individuals *after the dispersal step*.

These quantities are needed to explicitly compute the growth rate of the mutant in old-habitat patches, and to approximate the probability of adaptation. They are given by

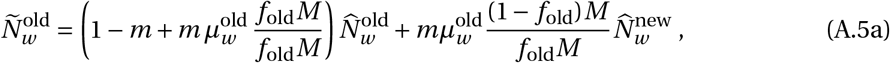

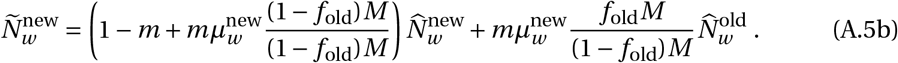

We then replace 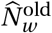 by *K*_o_l¿ (since old-habitat patches are assumed to be at carrying capacity after density regulation), and 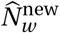 by the formula given in Eq. (A.4b). The shapes of 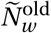 and 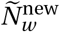 as functions of *m* are given in Fig. S2.

**Figure S2:**
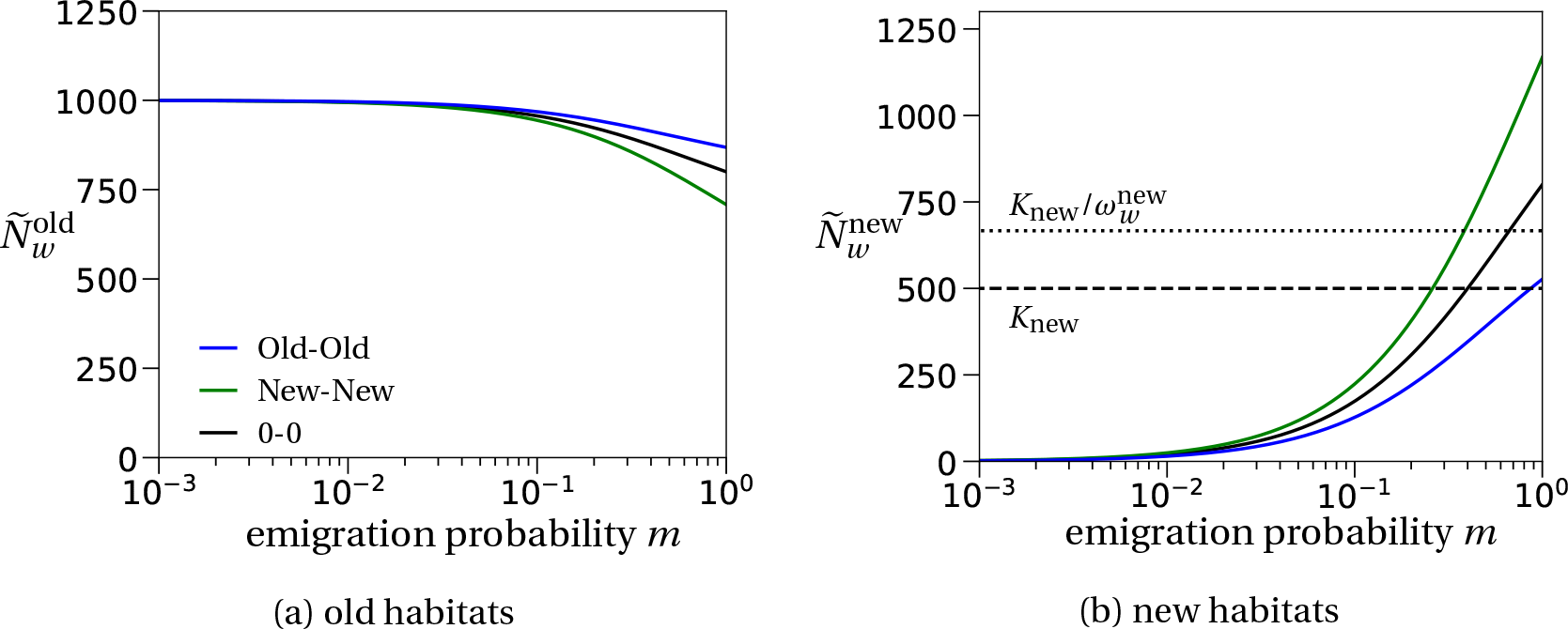
Wild-type population sizes after dispersal. As in Fig. S1, the wild-type population sizes after dispersal in (a) the old habitat and (b) the new habitat only depend on the wild-type immigration bias. We thus only plot the curves for the symmetric dispersal schemes. In (b) the dashed line is the reference value of the carrying capacity in new habitats, *K*_new_, and the dotted line represents the threshold from which on the carrying capacity will (on average) be exceeded after reproduction (see also Eq. (A.7) below). Parameters: Table 1 with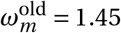.

### A.3. Local per capita growth rates

#### A.3.1. The local per capita growth rate *a*_old_

As stated in Eq. (2) in the main text, we define the per capita growth rate of rare mutant in the old habitat by

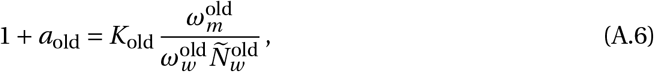

where we replace 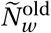 by the formula given in (A.5a).

#### A.3.2. The local per capita growth rate *a*_new_

In new-habitat patches, we may observe two situations after the reproduction step; either the carrying capacity is exceeded, in which case density regulation takes place, or the carrying capacity is not reached after reproduction. The former scenario occurs if the wild-type population size after dispersal, 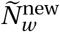, exceeds the threshold 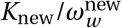. This threshold corresponds to the wild-type population size that will (on average) decrease to *K*_new_ after reproduction. In this case, the local per capita growth rate of the mutant takes the same form as in old-habitat patches. In contrast, if the carrying capacity *K*_new_ is not reached after reproduction, the local per capita growth rate of the mutant only depends on its fecundity 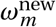. Combining these considerations gives the local per capita growth rate *a*_new_ as stated in Eq. (3) in the main text:

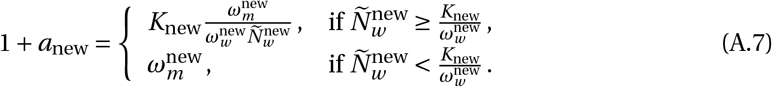

The shapes of the local per capita growth rates of the mutant as function of *m* are illustrated in Fig. S3.

**Figure S3:**
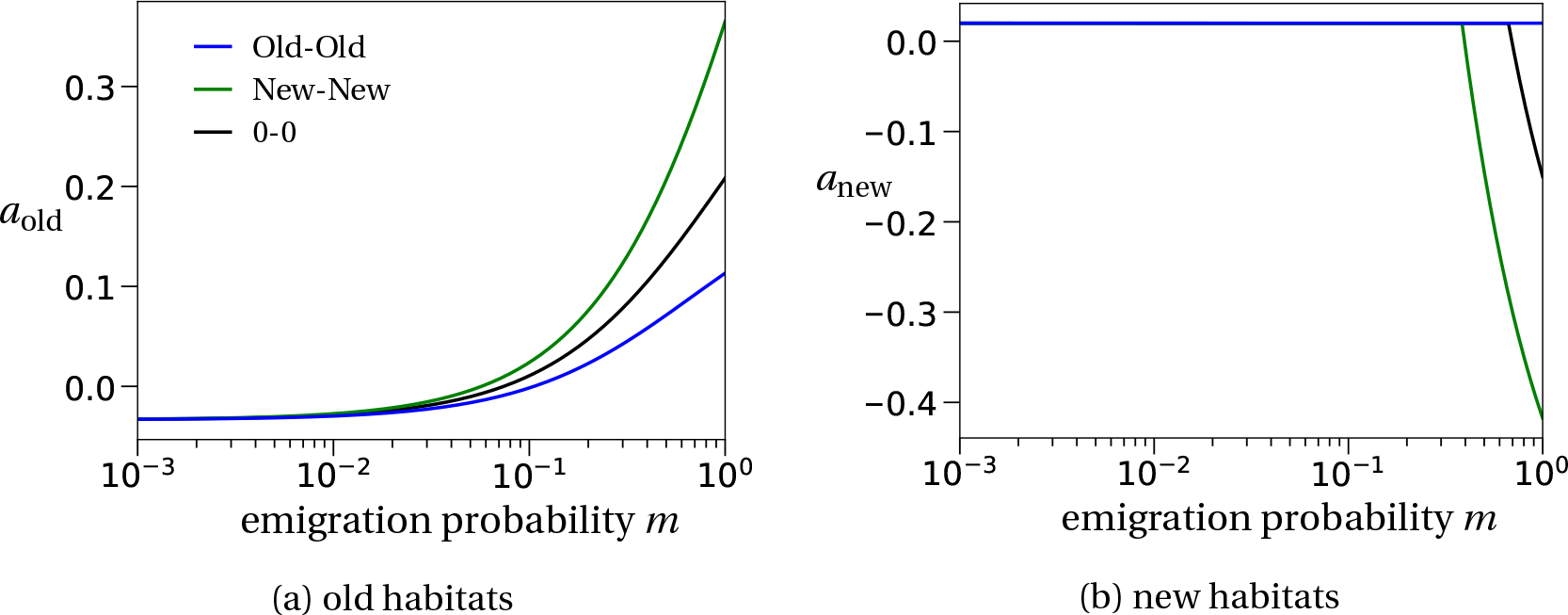
Local per capita growth rate of the mutant. (a) The local growth rate in old-habitat patches, *a*_old_ increases with the emigration probability *m*. This is due to the decreasing wild-type population size 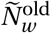 with increasing *m*. (b) The local growth rate in new habitats, *a*_new_, is barely affected by changes in *m*. We see a decrease in the growth rate only for very large emigration rates and dispersal schemes where the wild type does not preferentially immigrate into old-habitat patches. In this parameter range, the number of wild-type immigrants is so large and exceeds the threshold 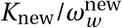, see Fig. S2b, that growth of better adapted mutant individuals is inhibited. Parameters are given in Table 1 with 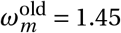.

### A.4. Wild-type population sizes during environmental change

Lastly, we compute the (deterministic) wild-type population size over time during the environmental change. This value is used in the approximation of the probability of evolutionary rescue in Eq. (12) in the main text, more precisely to estimate the number of rescue mutants that appear during the deterioration of patches.

At the moment a patch deteriorates, its population size is still given by the carrying capacity of the old habitat, *K*_old_. We order the patches in their sequence of deterioration, i.e., patch 1 deteriorates first and so on. Since all patches are connected to one another, this labelling has no effect on the dynamics. The number of wild-type individuals in patch *i* at time *t* is then given by

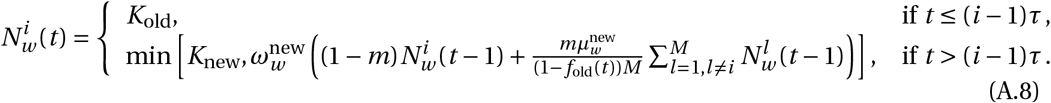

## B. Approximation of the establishment probability

We compute the survival probability of the lineage of a single mutant starting either in an old- or in a new-habitat patch. We call this probability the establishment probability because it implies the successful establishment of a mutant population within the metapopulation. It is denoted by *φ_k_, k* indicating the initial habitat type of the mutant (old or new).

Our method is the same as the one used in Tomasini and Peischl (2018), with the exception that our per capita growth rates of the mutant, *a*_old_ and *a*new, depend on the demography of the population. The method in general is based on the theory of multi-type branching processes, cf. Chapter 5.5 in Haccou et al. (2005). We refer the reader to the Supplementary Information of Tomasini and Peischl (2018) for a detailed application of the theory.

The mean reproduction matrix 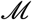 of a mutant gives the average number of offspring in a certain habitat, depending on the habitat type in which the mutant resides (see also Eq. (4) in the main text):

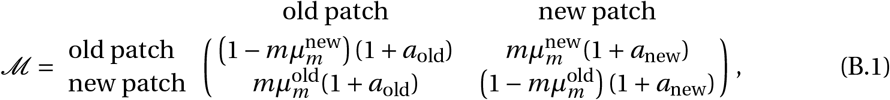

where the rows denote the parent locations, and the columns the patch type of the offspring.

Our goal is to apply Theorem 5.6 from Haccou et al. (2005) which states that for a slightly super-critical branching process, i.e. where the survival probability is slightly above zero, the establishment probability can be expressed in terms of the largest eigenvalue *ρ* and the corresponding left- and right-eigenvectors of the mean reproduction matrix 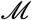, denoted by *u* and *v*, respectively. The eigenvectors should be normalized in the following way: *u*_1_ + *u*_2_ = 1 and 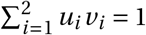. The establishment probabilities are then given by

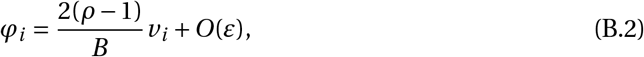

with

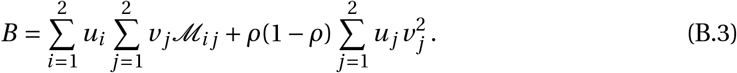

### B.1. Computing the largest eigenvalue

We first approximate the largest eigenvalue of 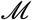 denoted by *ρ*. It is given by (see *Mathematica* notebook)

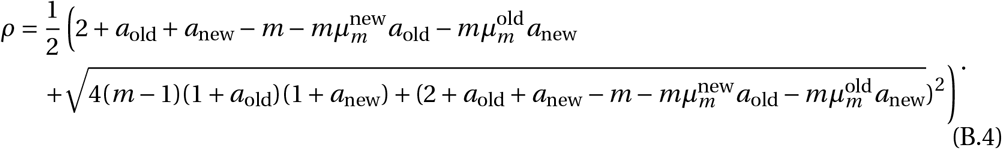

To make analytical progress and to identify under which conditions the process is slightly supercritical, i.e. *ρ >* 1, we rescale the parameters by a small parameter *ε*. We set *a*_old_ = *ε, a*_new_ = *ε* and *m = εη*. Assuming that *ε* is small enough, i.e. effectively a weak selection and weak dispersal assumption, we can neglect higher orders of *ε* and find

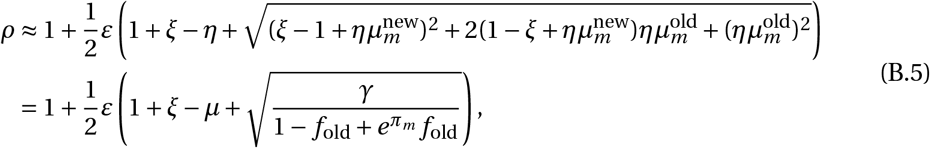

where *γ* is the rescaled version of the constant *C* in the main text (Eq. (8c)), i.e.

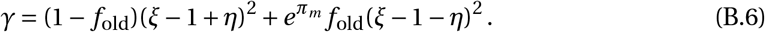

For *ε →* 0, we find that *ρ →* 1 (Eq. (B.5)) which means that the branching process is slightly super-critical if *ρ >* 1 and real. A sufficient condition for this to be true is that the first term in the parentheses of (B.5) is positive:

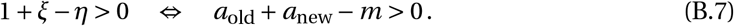

In the case that the branching process is not super-critical, the expression in Eq. (B.2) becomes negative and the establishment probability is zero.

### B.2. Computing the establishment probability

Eq. (B.2) involves normalized eigenvectors. Their precise form is of not much insight. We therefore omit stating them explicitly but refer to the *Mathematica* notebook. Solving Eq. (B.2) to the first order of *ε* and transforming back to the original variables, we obtain, after reordering, the equation presented in the main text (Eq. (8)).

## C. Disappearance of the relaxed competition effect

As we have argued in the main text, the increase of the establishment probability for large emigration probabilities *m* is due to the effect of relaxed competition. More precisely, the local per capita growth rate of the mutant, *a*_old_ becomes positive. If the fecundity of the mutant in the old habitat, 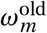 is too low, we do not observe relaxed competition in old habitats and therefore no increase of the establishment probability for large emigration probabilities *m* (Fig. S4). For the dispersal schemes New-Old and New-New, relaxed competition is still visible for very large values of *m*, which is explained by the strong immigration bias of wild-type individuals to the new habitat (see also Fig. S2).

**Figure S4:**
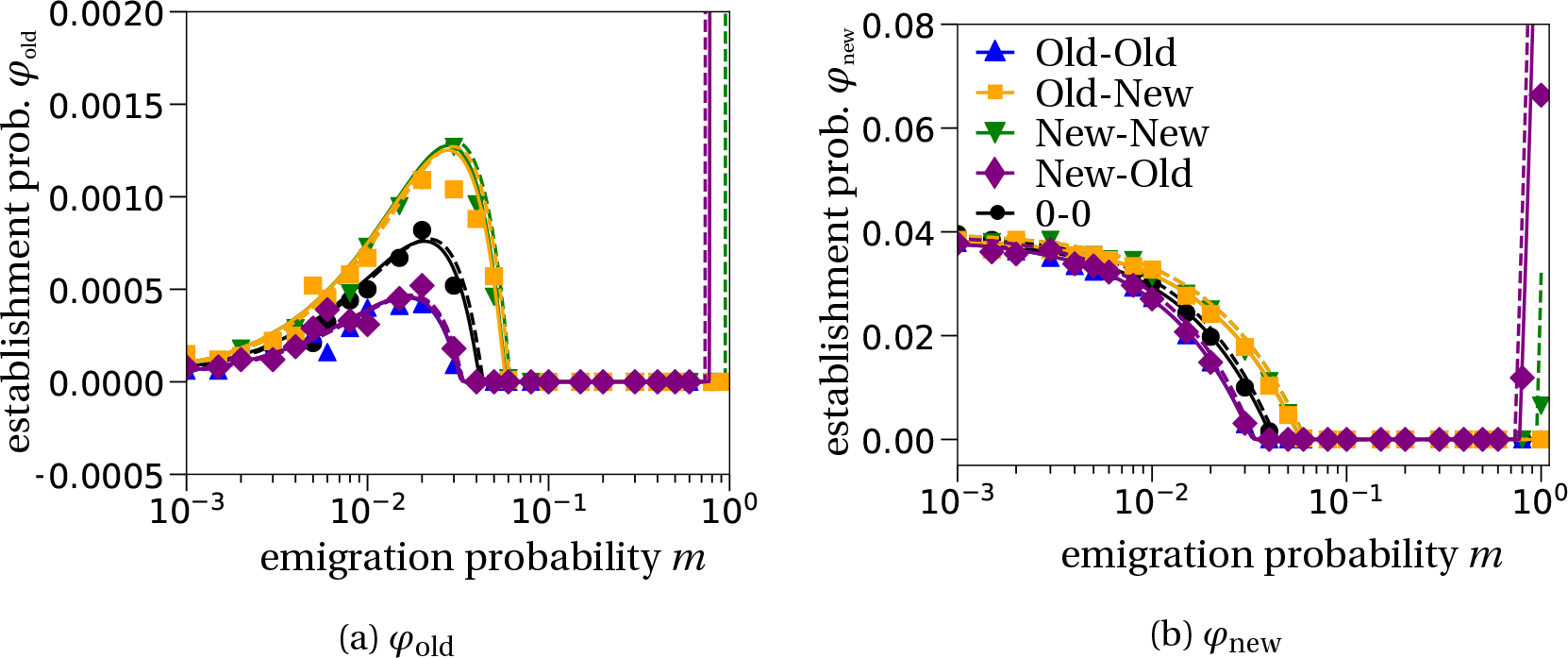
Disappearance of region (iii) for large fecundity differences in the old habitat. If the mutant fecundity in old-habitat patches is too low, here 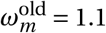, the effect of relaxed competition is not strong enough to have an impact on the establishment probability for high dispersal rates unless there are very few wild-type individuals left in old-habitat patches after dispersal. This can only happen for a wild-type dispersal bias towards the new habitat, i.e. the New-New and New-Old dispersal schemes. The establishment probability for the other dispersal schemes remains at zero.

## D. Habitat of origin of the adaptive mutation

Fig. 4 illustrates that most successful adaptive mutations arise in old-habitat patches. Here, we disentangle the two factors that contribute to the emergence and establishment of mutants; the establishment probability *φ_k_* and the wild-type stationary population sizes. Fig. S5b shows that the large number of successful mutant lineages appearing in old-habitat patches is due to the large mutational input resulting from the much larger wild-type population sizes in old habitats (dashed line), compared to the wild-type population size in new habitats (solid line). The establishment probability alone, Fig. S5a, would predict more successful mutant lineages arising in new-habitat patches than in old-habitat patches, i.e., *φ*_new_ is always larger than *φ*_old_.

**Figure S5:**
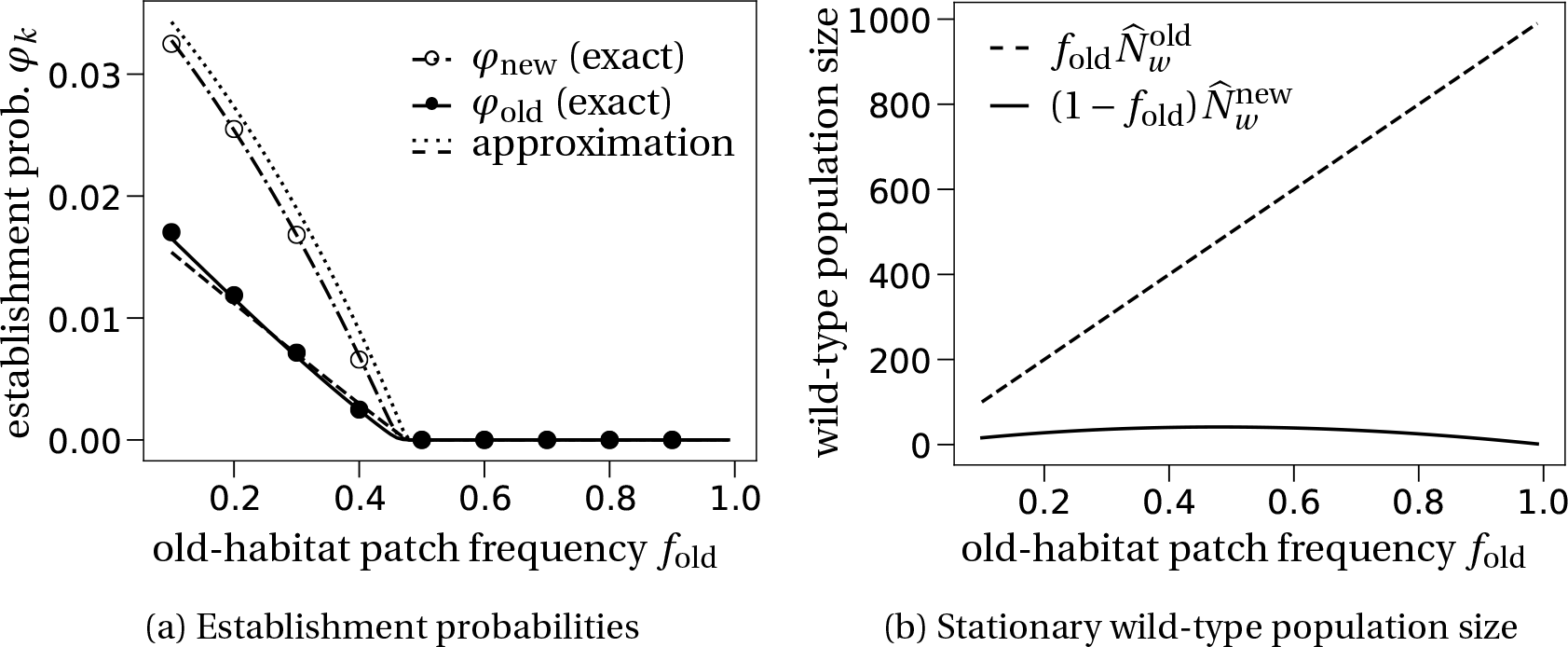
Establishment probability and stationary wild-type population size when varying the old-habitat frequency. In the simulations we have used the standard set of parameters as given in Table 1 and the unbiased dispersal scheme (*π_w_ = π_m_* = 0). In (a) we additionally chose the large fecundity difference scenario 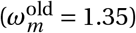.

## E. The speed of environmental degradation and standing genetic variation

We have seen in Fig. 6b that standing genetic variation can have a non-negligible part in the process of evolutionary rescue. Here, we study how the contribution of standing genetic variation changes with the speed of the degradation process, i.e. as a function of the time between two degradation events *τ*. We find that if the time between two consecutive degradation events is larger than three generations (*τ* > 3), the contribution of standing genetic variation to the probability of evolutionary rescue is smaller than the contribution by de-novo mutations, as illustrated in Fig. S6. This is explained by the longer time span that mutants need to survive to experience a beneficial environmental configuration. For example, in the chosen parameter set, there need to be more than three degraded patches for the mutant to have a positive probability of establishment (see Fig. 3d). Therefore, smaller values of *τ* increase the probability that mutants that were present in the population before the first patch degraded survive up to this favorable environmental configuration.

**Figure S6:**
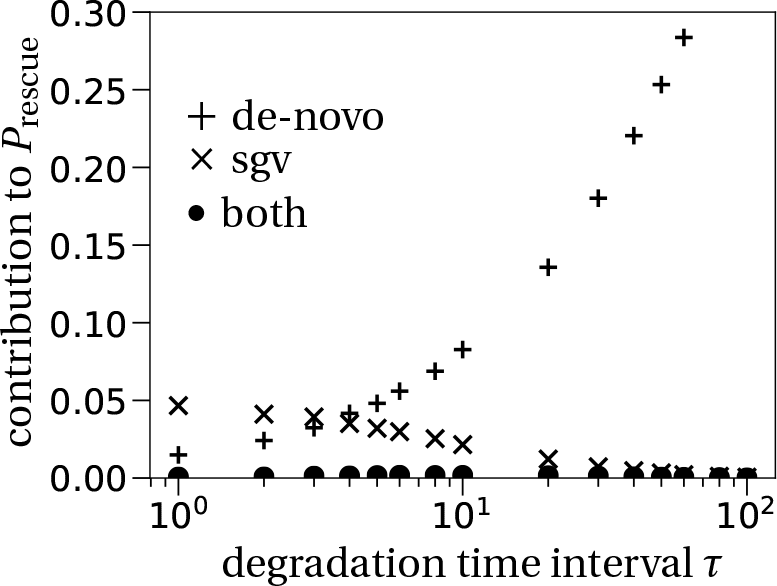
Disentangling contribution of standing genetic variation (sgv) and de-novo mutations to the probability of evolutionary rescue as a function of the speed of the environmental degradation process *τ*. With our parameters, if the time between two degradation events is longer than 3 generations, the contribution of rescue events due to de-novo mutants is larger than the contribution by standing genetic variation. Mutant individuals that are present at the first degradation event need to survive a deleterious environmental configuration where they have no chance of establishing a mutant subpopulation, cf. Fig. 3b,d. Fast environmental degradation (small *τ*) increases the probability that these mutants survive until a beneficial environmental configuration. Parameters: 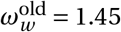, unbiased dispersal scheme (*π_w_ = π_m_* = 0), other parameters as stated in Table 1.

## F. Establishment probability in a model without demography

In this section, we consider a variation of our original model to investigate the impact of demography on the establishment probabilities *φ*_old_ and *φ*_new_. The dispersal process and the dynamics in old-habitat patches remain as studied before. We now assume that the population remains at carrying capacity in new-habitat patches, i.e. there is no longer a declining wildtype population, so that the stationary wild-type population size is 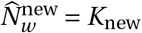. For simplicity we will also assume that *K*_new_ = *K*_old_ = *K*. Since there is now always density regulation in new habitat patches as in old-habitat patches (implemented by hypergeometric sampling as before), the per capita growth rate of the mutant in new-habitat patches is given by

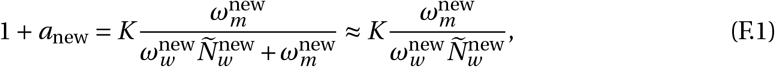

where we replace 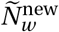 by the formula provided in Eq. (A.5b), with 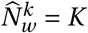. The rest of the mathematical analysis remains unchanged and as described before (Sections A and B).

We see that, as mentioned in the main text, region (iii) of the establishment probability disappears in these type of models except for the Old-New and the New-Old dispersal schemes. The reason for the disappearance of the region is that relaxed competition only plays a subordinate role for the type-independent dispersal schemes (Old-Old, New-New and 0-0). In other words, these dispersal schemes maintain the local frequencies of the mutant at the same level as before the dispersal step and by that do not change the population dynamics. In contrast, the type-dependent dispersal schemes, Old-New and New-Old, distribute the wild type and the mutant into separate patch types. For large emigration probabilities *m*, this results in relaxed competition for mutant individuals, hence explaining the increase of establishment probability for large values of *m*.

**Figure S7:**
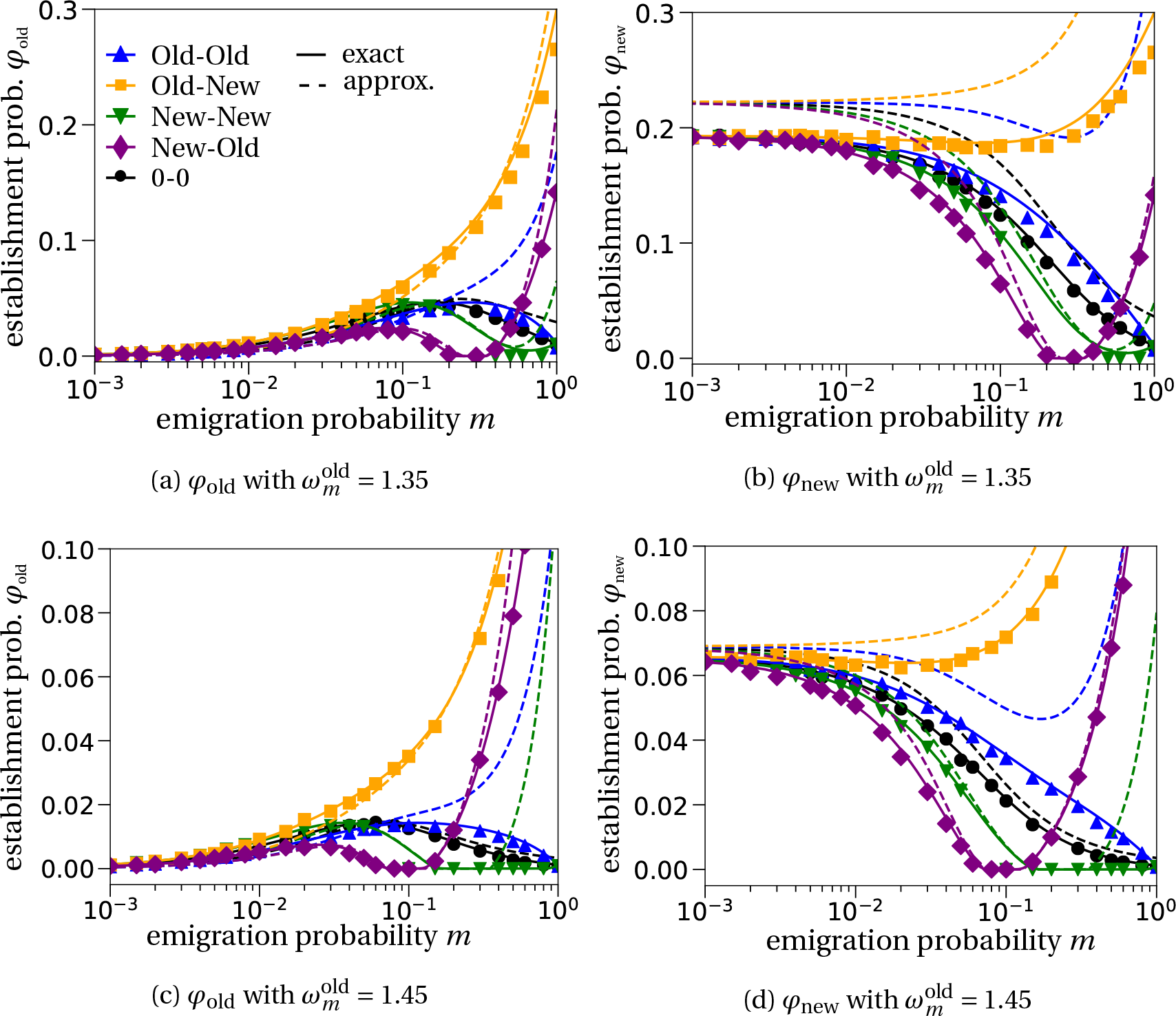
Establishment probability when populations in both habitats are at carrying capacity. We plot the establishment probability for a single mutant either initially in an old-habitat patch (a,c) or in a new-habitat patch (b,d). The numerical solution (solid lines) still approximates the simulated data reasonably well. The analytical approximation (dashed lines) however deviates strongly from the data due to large growth rates (*a*_new_ ≈ 0.2) so that the conditions for the approximation to hold are violated. In this case, in Eq. (8) higher order corrections would need to be taken into account. The fecundity values in the new habitat are given by 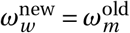 and 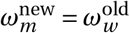 and the carrying capacity is *K*_new_ = *K*_old_ = 500. The other parameters are as stated in Table 1. Missing data points (mostly for the negative density-dependent dispersal scheme – green triangles) are explained by too large computation times. All data points are averages from 10^4^ independent runs.

## G. Habitat of origin depending on the dispersal scheme

The habitat type of the origin of the rescue mutation is depends on the considered dispersal scheme. Fig. S8 shows the relative contribution of each natal habitat type to the probability of evolutionary rescue (with 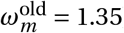). The establishment probabilities are close among the three type-independent dispersal schemes (0-0, Old-Old, and New-New). For intermediate emigration rates *m*, the Old-New dispersal scheme shows an excess, when compared to the other dispersal schemes, in successful mutant lineages arising in old habitats. As *m* increases, this difference between the dispersal schemes diminishes. For large *m*, the New-Old dispersal scheme displays a strong contribution of mutants emerging from old-habitat patches. A possible explanation is the large probability of establishment for a mutant emerging in old-habitat patches for this parameter set (cf. Fig. 2a).

**Figure S8:**
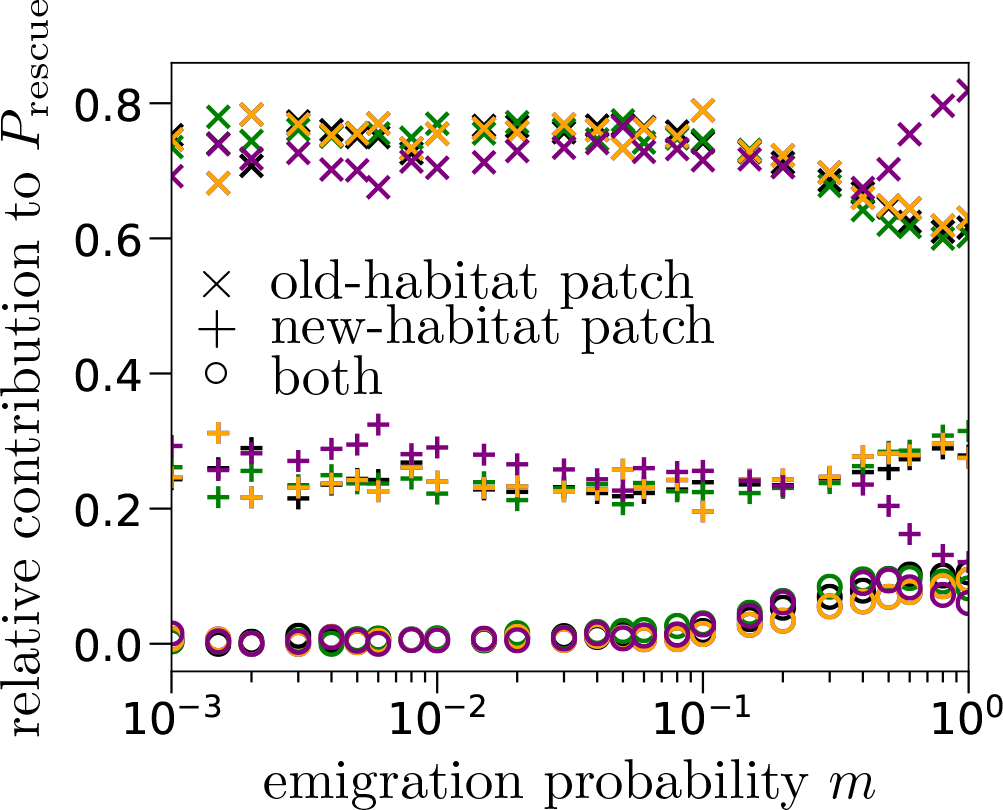
Habitat type of the origin of the rescue mutant depending on the dispersal scheme. Relative contributions of each habitat type to the probability of evolutionary rescue, as a function of the emigration probability *m*. The color-coding is as in the main text: black for the 0-0, blue for the Old-Old, green for the New-New, orange for the Old-New, and purple for the New-Old dispersal scheme.

## H. Large mutant fecundity in the new habitat

In the main text, we have assumed that the mutant fecundity in the new habitat, 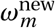, is just slightly above 1. Here, we show that increasing this value will simply increase the probability of establishment and of evolutionary rescue, as illustrated in Figs. S9.

For the establishment probability, we observe a similar pattern as in Fig. 2c,d. If the mutant individual appears in the old habitat, the establishment probability *φ*_old_ is a strictly increasing function of the emigration probability *m*. The only exception is the New-New dispersal scheme where gene swamping can occur for large emigration probabilities *m*. The New-Old dispersal scheme can compensate the gene swamping effect by relaxed competition in the old habitat. The parameter set we plot in Fig. S9 corresponds to the one used to plot Figs. 2a,b. Comparing these subfigures with each other, we see that the local maximum for intermediate emigration probabilities *m* is not visible for a larger mutant fecundity 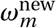. Additionally, the change in mutant fecundity in the new habitat 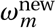 affects the quantitative results of the establishment probability. The shape of the establishment probability can be explained by the same means as illustrated in the main text.

**Figure S9:**
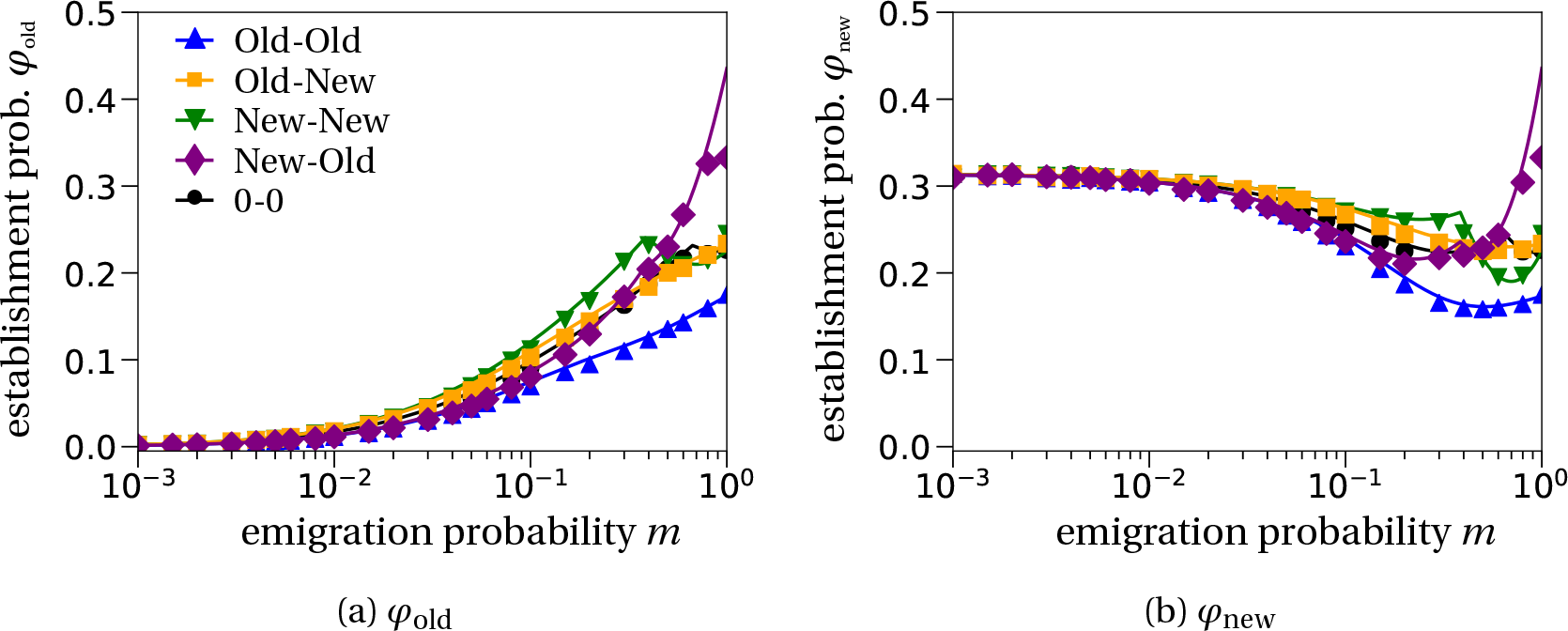
Establishment probabilities for large mutant fecundity in the new habitat. The mutant fecundities in the old and new habitat are set to 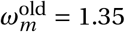 and 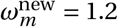, respectively. The other parameters are as given in Table 1. We only plot the numerical solution of Eq. (7) because the assumption of slight super-criticality of the branching process is violated here. Overall, we find a similar qualitative behavior as in Fig. 2 in the main text. The initial increase in *φ*_old_ (decrease in *φ*_new_) is explained by a beneficial (deleterious) effect of the emigration probability on the change of habitat of the mutant individual. The non-monotonic behavior for large values of *m* is a combination of gene swamping, which reduces the establishment probability, and relaxed competition, which increases the establishment probability.

## Footnotes

1 https://gitlab.com/pczuppon/evolutionary_rescue_and_dispersal

2 https://gitlab.com/pczuppon/evolutionary_rescue_and_dispersal

